# Molecular anatomy of the glomerular filtration barrier reveals topology-encoded albumin sieving

**DOI:** 10.64898/2026.09.30.755551

**Authors:** Xing Wang, Weiyan Zheng, Wenjing Du, Dongxuan Chi, Wenjia Fan, Ruoheng Mo, Jie Ding, Li Yang, Jing Nie, Ming Han, Qiang Guo

## Abstract

The kidney filters plasma rapidly while retaining albumin, yet where and how this selectivity is encoded within the glomerular filtration barrier remain unresolved. Here we combine cryo-electron tomography (cryo-ET) of vitrified mouse glomeruli with structure-resolved transport modelling to predict albumin passage from the native three-dimensional architecture of the barrier. Among the three barrier layers, endothelial fenestrae and the flexible Nephrin-Neph1 slit-diaphragm lattice are substantially permeable to albumin. The glomerular basement membrane (GBM) instead provides the dominant restriction through steric exclusion governed by void-network topology. In local GBM reconstructions, water traverses broadly connected void space, whereas narrow throats exclude albumin, leaving few accessible paths. To scale this mechanism to the tissue level, we combine structural statistics from local cryo-ET volumes with mesoscale measurements of GBM heterogeneity to generate representative ensembles. Integrating these ensembles with other layers predicts whole-barrier albumin sieving in close agreement with an independent measurement. The topology-driven mechanism also explains early filtration failure in Alport syndrome mice, in which collagen-IV disruption remodels the GBM network and opens albumin-accessible paths while cellular interfaces remain preserved. These findings identify the structural origin of glomerular albumin selectivity and establish a quantitative route from locally resolved architecture to tissue-scale physiology.

## Introduction

Structural biology derives biological function from molecular architecture, but extending this principle from macromolecular assemblies to tissue physiology remains a major challenge. At the tissue scale, function emerges from heterogeneous structures spanning cellular and extracellular compartments, with interactions extending across multiple length scales. The glomerular filtration barrier (GFB) is a multiscale tissue system central to kidney function, sustaining rapid plasma filtration while retaining albumin and other circulating macromolecules. This selective function is distributed across a fenestrated endothelium, the glomerular basement membrane (GBM), and podocyte foot processes connected by slit diaphragms ^1–3^. Failure of this barrier causes proteinuria and progressive kidney disease, yet the native structural basis of albumin selectivity remains unresolved.

Which GFB layer dominates albumin restriction, and through what structural mechanism, remains debated. Genetic perturbations and disease phenotypes establish the functional necessity of both the GBM and slit diaphragm. GBM disruption is a hallmark of collagen-IV disorders such as Alport syndrome ^4–7^. Loss of the slit-diaphragm proteins Nephrin or Neph1 causes severe proteinuria, supporting the long-standing view that the slit diaphragm forms the terminal molecular sieve ^8–11^. Recent structural studies reveal flexible, multilayered or fishnet-like slit-diaphragm assemblies, establishing the structural foundation for investigating its molecular organization and filtration properties ^12–15^. However, whether this architecture provides sufficient resistance to explain albumin retention across the intact multilayer filtration barrier has remained unresolved. Physiological models infer effective size- and charge-selective properties from bulk transport measurements, but often assume idealized local geometries that cannot capture the structural heterogeneity of individual layers ^1,16^. Determining how albumin-selective resistance is distributed across the GFB therefore requires native, layer-resolved structures to be incorporated into a physical transport framework^17–19^.

Here, we combine cryo-electron tomography (cryo-ET) of vitrified mouse glomeruli with structure-resolved transport modeling to derive albumin transport from the molecular anatomy of the GFB. Endothelial fenestrae and the native slit-diaphragm lattice are substantially permeable, whereas sparse continuous paths through the GBM void network impose the dominant restriction. Serial integration of the three layers predicts albumin sieving in close agreement with independent measurements in perfused kidneys. In *Col4a3*-deficient mice, collagen-IV disruption reorganizes albumin-accessible GBM pathways while the cellular interfaces remain morphologically recognizable. These findings identify the GBM as the principal albumin-restrictive layer and establish a route from tissue cryo-ET to quantitative physiological prediction.

## Results

### Cryo-ET resolves the molecular organization of the intact glomerular filtration barrier

To establish a structural basis for layer-resolved filtration analysis, we developed a workflow that preserves isolated mouse glomeruli in a vitrified, hydrated state. Size-enriched glomeruli retained capillary-loop and podocyte organization ^20^, as assessed by immunostaining and conventional electron microscopy (Extended Data Fig. 1). Cryo-volume electron microscopy confirmed preservation of the filtration barrier at the mesoscale, resolving fenestrated endothelium, the GBM, and interdigitating podocyte foot processes across capillary loops (Fig. 1A–C, Supplymentary Movie 1). An optimized Cryo-focused ion beam (cryo-FIB) serial lift-out workflow ^21^ was then used to prepare lamellae containing the complete local filtration pathway for cryo-ET analysis.

**Fig. 1.**
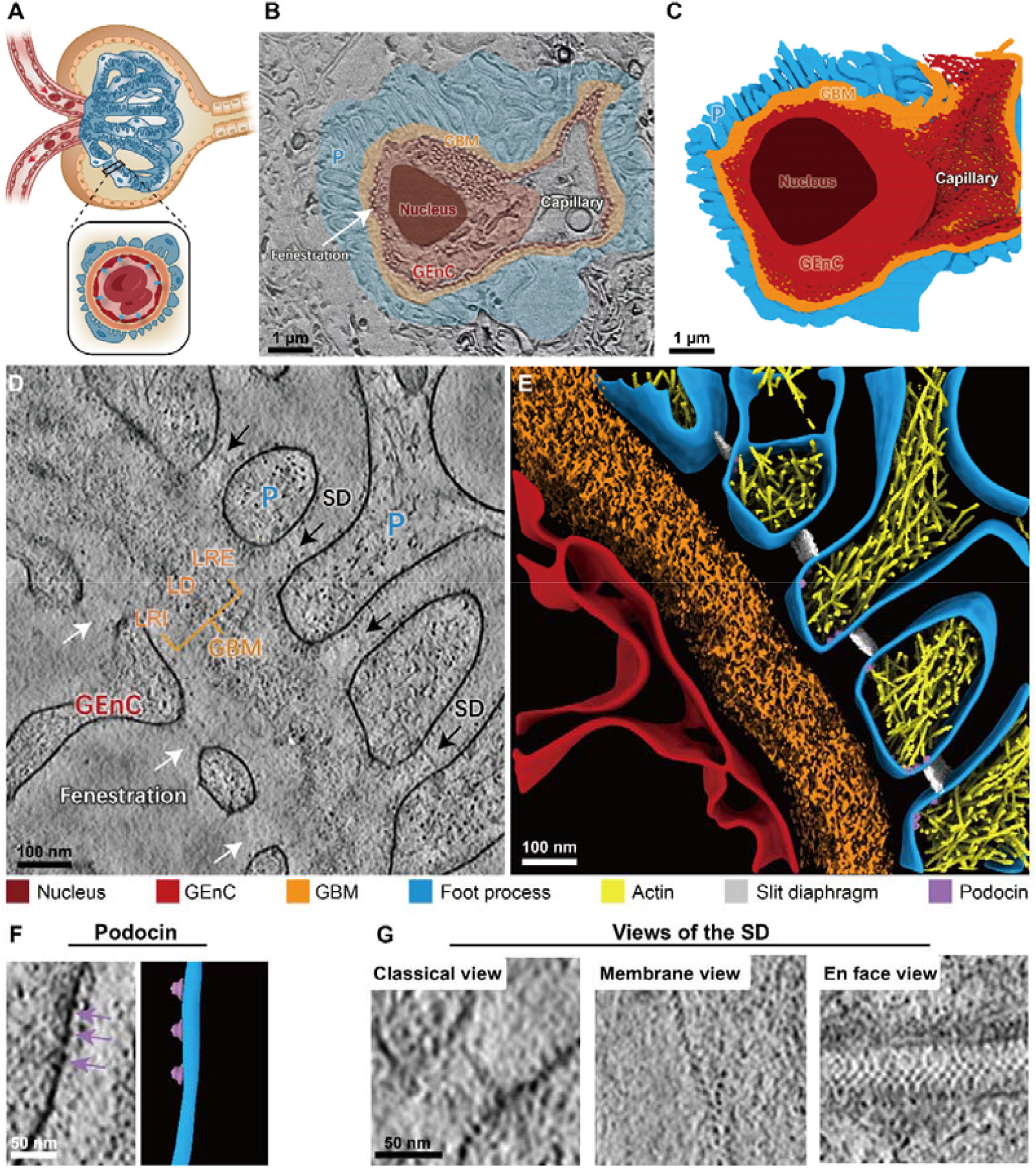
Multiscale architecture of glomerular filtration barrier. (A) Schematic of the glomerular microanatomy. The glomerulus consists of a capillary tuft connected to afferent and efferent arterioles, encapsulated by Bowman’s capsule. The boxed inset highlights the trilaminar glomerular filtration barrier (GFB), comprising the fenestrated endothelium (GEnC), glomerular basement membrane (GBM), and podocyte foot processes. (B–C) Three-dimensional characterization of the GFB by cryo-PFIB/SEM. (B) A representative image slice and (C) its corresponding 3D segmentation. The rendered volume was reconstructed from 83 serial slices acquired at 20 nm intervals, covering a total tissue volume of ~27.8 μm^3^. (D–E) Local ultrastructural characterization of the GFB by cryo-ET. (D) A representative tomographic slice visualized as a 10-slice-Z-projection (9.48 nm), alongside (E) its corresponding 3D segmentation of the local GFB region. (F) A representative tomographic slice (left) and corresponding 3D segmentation (right) of podocyte membrane and associated cage-like densities. Putative podocin densities are indicated by purple arrows. (G) Orthogonal views of slit diaphragm: sheet-like structure (left) in cross-section, periodic punctate densities in membrane view (middle), and a mesh-like arrangement in the en face view (right). Scale bars: 1 µm in B and C; 100 nm in D and E; 50 nm in F and G.

We collected 549 tomograms from seven glomeruli and resolved all three GFB layers within a shared near-native tissue context (Fig. 1D–E, Supplymentary Movie 2). Attenuated endothelial regions were interrupted by fenestrae that generally lacked detectable diaphragm-like densities at the current resolution, consistent with mature glomerular endothelium ^22^. The intervening GBM contained an electron-dense lamina densa composed of short, interwoven fibrillar elements (Fig. 1D–E). On the urinary side, podocyte foot processes contained dense actin bundles and were connected by a mesh-like slit diaphragm, whereas their plasma membranes displayed cage-like densities consistent with podocin assemblies (Fig. 1F–G). Together, these observations established a structural framework for determining how each GFB layer constrains albumin passage.

### Endothelial fenestrae form a geometrically permissive filtration interface

Filtration begins at the fenestrated endothelium. En face views showed quasi-circular fenestrae distributed across attenuated endothelial regions (Fig. 2A, Supplymentary Movie 3), with a mean diameter of approximately 91.3 nm and most openings spanning 60–120 nm. These dimensions greatly exceed those of albumin and most soluble plasma proteins but remain far smaller than circulating blood cells, indicating that the endothelium imposes little geometric restriction on albumin while excluding cellular components.

**Fig. 2.**
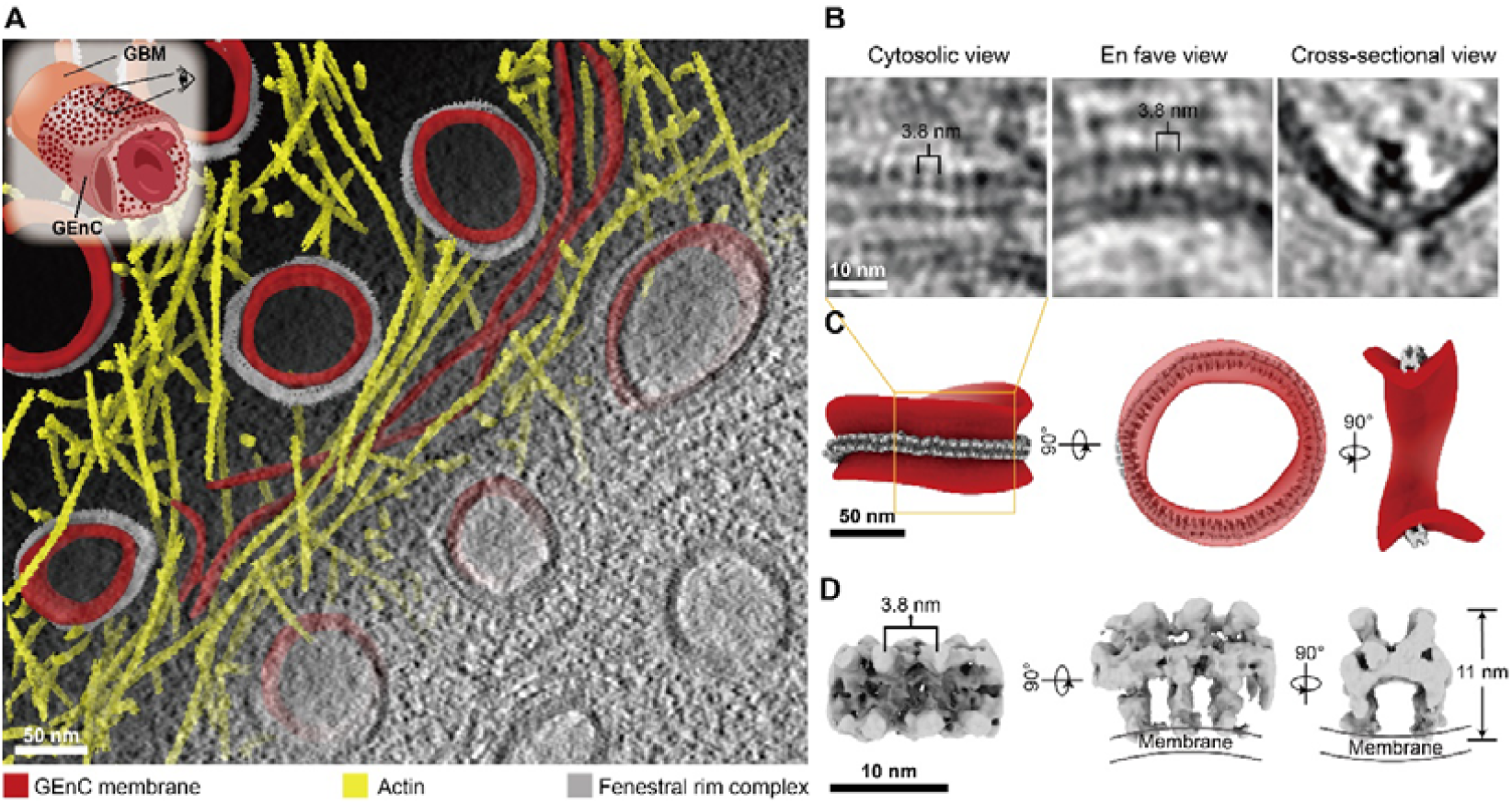
Cryo-ET reveals the ultrastructure of endothelial fenestrae. (A) En face tomographic slice of an endothelial cell (GEnC) region enriched with fenestrae, overlaid with the corresponding 3D segmentation using a gradient transition. Orientation is indicated in the inset. Densely distributed endothelial fenestrae are visible, lighter regions correspond to fenestral lumina/extracellular space. (B) Representative views of the fenestral region. Cytosolic and en face view reveal protein densities with 3.8 nm periodicity, whereas the cross-sectional view illustrates the fenestral rim complex in profile. (C) 3D surface rendering of the fenestration corresponding to the one shown in (B), with the plasma membrane colored in red and fenestral rim complex in grey. (D) Subtomogram average of the fenestral rim complex at 11 Å resolution. Scale bars: 50 nm in A and C; 10 nm in B and D.

Such functional geometry of endothelial fenestrae was molecularly organized. Cryo-ET revealed a periodic density that followed the highly curved cytoplasmic surface of each fenestral rim and formed a ring around the opening (Fig. 2B–C). Subtomogram averaging of 3,004 particles from 255 rim segments across 26 tomograms yielded an 11 Å reconstruction with a repeat spacing of approximately 3.8 nm (Fig. 2D and Extended Data Fig. 2). The rim complex resembled membrane-bound EHD-family filaments in both periodicity and curvature-dependent localization^23,24^, and the enrichment of EHD3 in glomerular endothelium further supports this putative assignment ^25–27^. The assembly may stabilize fenestral curvature while preserving a pore size suited to high-throughput plasma filtration.

### Native slit-diaphragm architecture permits substantial albumin passage

The slit diaphragm has long been considered a leading candidate for the albumin-selective barrier because disruption of its core proteins causes severe proteinuria ^8,9^. Consistent with previous cryo-ET studies, *en face* views of our raw tomograms showed the characteristic fishnet-like morphology of the slit diaphragm ^13–15^. Subtomogram averaging resolved its extracellular assembly at approximately 20 Å resolution (Extended Data Fig. 3). The density formed a non-polarized periodic lattice spanning adjacent foot processes, with crisscrossing strands bridging the slit at approximately 12-nm intervals. Triangular membrane-proximal densities linked the extracellular lattice to the podocyte surface, thereby defining the membrane-associated organization of the fishnet-like lattice in the subtomogram average (Fig. 3A and Extended Data Fig. 4).

**Fig. 3.**
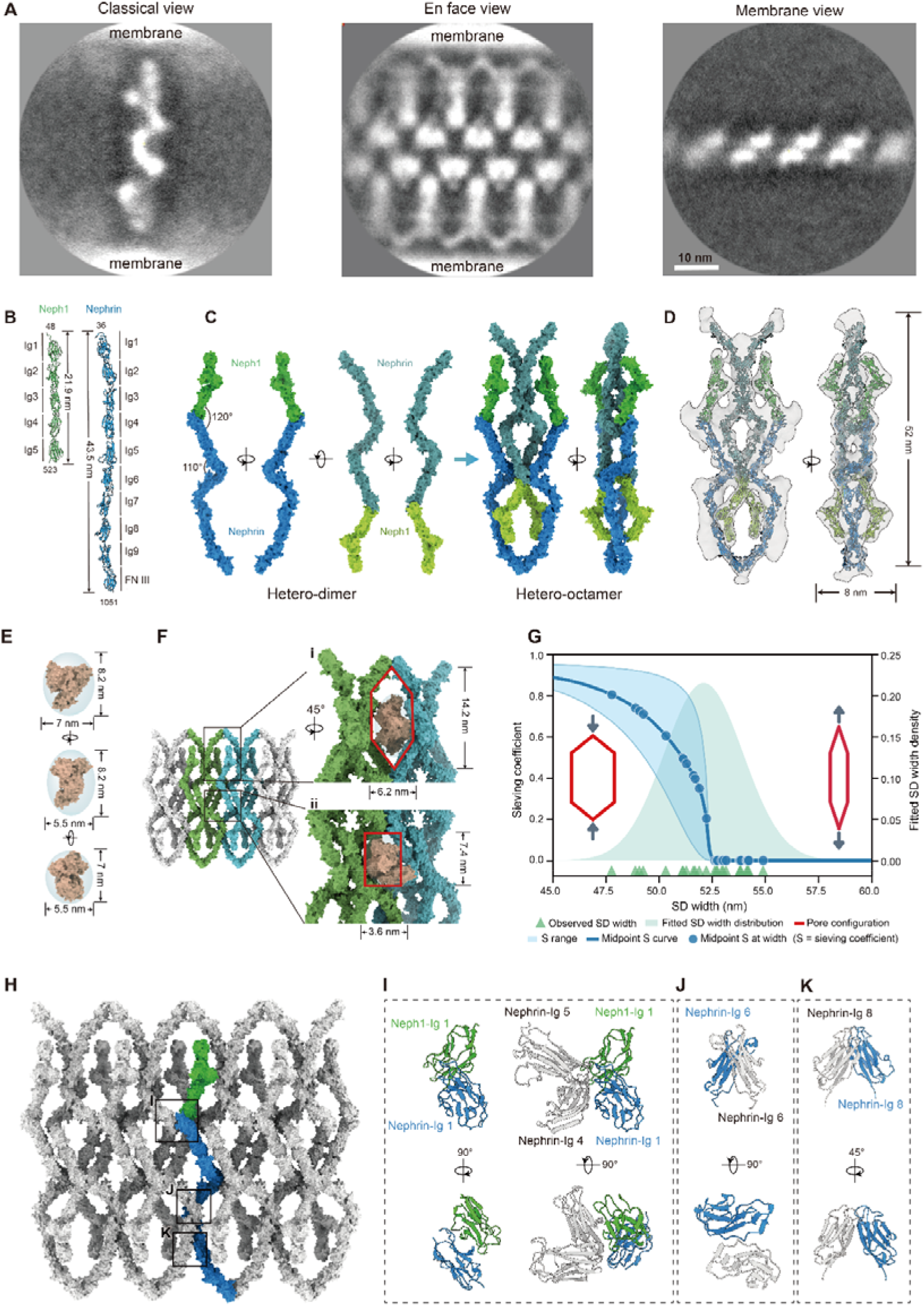
Molecular architecture of the slit diaphragm. (A) Slit diaphragm architecture resolved by subtomogram averaging, shown in three orientations: classical cross-sectional view perpendicular to the filtration axis, en face view parallel to the filtration axis, and membrane view perpendicular to the podocyte membrane. The periodicity of the slit diaphragm was measured to be 12 nm in both the en face and membrane views. Scale bars: 10 nm. (B) Domain organization of Nephrin and Neph1 ectodomains. Nephrin (residues 36–1051, AF-Q9QZS7-F1-v6) comprises nine N-terminal immunoglobulin-like C2 domains (Ig1–Ig9) and a C-terminal fibronectin type III domain (FNIII). Neph1 residues (48–523, AF-Q80W68-F1-v6) contains five N-terminal Ig-like C2 domains (Ig1–Ig5). (C) Molecular model of the Nephrin-Neph1 complex. Nephrin (blue) and Neph1 (green) form an inter-molecular angle of ~120°, with a ~110° bend at the Nephrin Ig4–Ig5 interface. The hetero-octameric assembly is generated via rotation and translation; vertically flipped heterodimers are indicated in lighter shades (Nephrin, cadet blue; Neph1, green yellow). (D) Fitting of the slit diaphragm molecular model into the subtomogram averaged density map, shown in two orthogonal orientations. (E) Geometric approximation of albumin as a triaxial ellipsoid. The molecular structure of albumin (PDB:1AO6) is modeled as an ellipsoid with dimensions of approximately 8.2 nm × 7 nm × 5.5 nm. Three different projection views show the corresponding semi-axes and spatial envelopes. (F) Geometric approximation of the slit diaphragm lattice openings. The model identifies two types of channels: a large channel with major and minor axes of approximately 14.2 × 6.2 nm, permeable to albumin; and a small channel of approximately 7.4 × 3.6 nm, predicted to be sterically restrictive to albumin. (G) Predicted albumin sieving coefficient across slit diaphragm width-dependent pore deformation using orientation-bounded accessibility estimates. The fitted slit diaphragm width distribution is shown as the shaded cyan area, and observed slit diaphragm width measurements are indicated by teal triangles. The red hexagons schematically represent pore configurations corresponding to the minimum (left) and maximum (right) slit diaphragm stretching. The light-blue band shows the range of predicted sieving coefficients bounded by random three-dimensional albumin orientation and best-passage orientation, and the blue curve shows the midpoint of this lower-upper range. Blue circles indicate midpoint sieving coefficient values at observed slit diaphragm widths. denotes the albumin sieving coefficient. The integrated midpoint slit diaphragm sieving coefficient calculated from the fitted SD width distribution is 0.48. (H) Idealized model of the slit diaphragm assembly. The slit diaphragm is composed of four hetero-octamers, with a representative Nephrin (blue)-Neph1 (green) heterodimer highlighted. The black box outlines potential interaction regions. (I) Primary slit diaphragm interaction interface. The assembly is stabilized by *trans*-heterophilic binding between Nephrin-Ig1 (blue) and Neph1-Ig1 (green), and *trans*-homophilic interactions between adjacent Nephrin-Ig4 and Nephrin-Ig5 domains (white). (J) Nephrin-Nephrin *cis*-homophilic interface, mediated by Nephrin-Ig6 (blue and white). (K) Membrane-proximal Nephrin-Nephrin interaction, where Nephrin-Ig8 (blue) engages in *cis*-homophilic binding with an adjacent Nephrin-Ig8 (white) near the podocyte membrane.

To define the molecular organization underlying the native lattice, we constructed an integrative model constrained by the cryo-ET density, sequence connectivity, published stoichiometry ^13^ and domain-interaction data ^28^. In the model, Nephrin and Neph1 formed a curved hetero-octameric assembly approximately 52 nm wide and 8 nm thick (Fig. 3B–D and Extended Data Fig. 5A–B, Supplymentary Movie 4). The assembly contained large apertures of approximately 14.2 × 6.2 nm and smaller apertures of approximately 7.4 × 3.6 nm (Fig. 3E–F). Albumin, with approximate dimensions of 5.5 × 7.0 × 8.2 nm (PDB: 1AO6), was excluded from the small apertures but lay near the steric limit of the large ones, permitting passage only in favourable orientations.

The native lattice, however, is not rigid: the distance between opposing foot-process membranes varied from 48 to 55 nm apart (Fig. 3G). Changes in slit diaphragm width did not uniformly enlarge or constrict the lattice. Narrower configurations compressed the pore across the slit while expanding the lateral dimension that limits albumin passage, thereby increasing the range of orientations compatible with transit. Sampling pore geometries across the measured width distribution yielded an integrated slit diaphragm sieving coefficient of approximately 0.48 (Fig. 3G). The slit diaphragm therefore imposes finite resistance and cannot account for the magnitude of albumin retention across the intact GFB.

Then what is the major role of slit diaphragm? Interface analysis of the integrative model predicted that the lattice is stabilized by heterophilic Nephrin-Neph1 interactions together with multiple homotypic Nephrin-Nephrin contacts (Fig. 3H). These include trans contacts between Nephrins on opposing foot processes, mediated by Ig1-Ig4/5 interactions, and cis contacts between Nephrins on the same foot process, involving the Ig6 and Ig8 domains (Fig. 3I–K). By linking each Nephrin-Neph1 unit to adjacent units, these interactions provide a structural basis for a continuous yet flexible adhesive lattice.

Disease-associated variants provided an independent test of the model-predicted interfaces. We mapped pathogenic variants in *NPHS1* and *KIRREL1*, which encode Nephrin and Neph1, respectively ^29^. Because *KIRREL1* variants predominantly impair plasma-membrane trafficking ^30^, we focused on extracellular *NPHS1* variants that could directly affect the predicted lattice interfaces. Nine pathogenic missense variants mapped to Nephrin-Nephrin contact surfaces across the Ig4, Ig5 and Ig8 domains (Extended Data Fig. 5C–F). Four of these variants, mL390V, mR422Q, mR757C and mV836M, have been reported to reach the cell surface ^31,32^. Their positions suggests that impaired extracellular assembly rather than defective trafficking alone, may contribute to disease. This provids a structural rationale for the essential role of slit-diaphragm proteins without requiring the lattice to function as a stringent albumin sieve.

### The glomerular basement membrane forms a mesoporous matrix that restricts albumin transport

Neither the endothelium nor the slit diaphragm could account for albumin retention, suggesting that the intervening GBM is the dominant restrictive layer. Cryo-ET resolved the electron-dense central region of the lamina densa as a connected meshwork of short fibrils approximately 3 nm in diameter (Fig. 4A–B, D), consistent with type IV collagen and laminin polymers^33,34^. By contrast, the flanking low-density regions contain substantially fewer fibrils therefore contribute little to albumin retention. Because stable adsorption to a finite number of matrix sites cannot sustain albumin retention during continuous filtration, we modelled the lamina densa as a restrictive porous medium. In this framework, albumin passage is governed by the accessibility and connectivity of inter-fibrillar voids. We segmented the lamina densa in 20 representative tomograms and represented the void space in each as a pore-throat network, with pores corresponding to local cavities and throats to the channels connecting them (Fig. 4B).

**Fig. 4.**
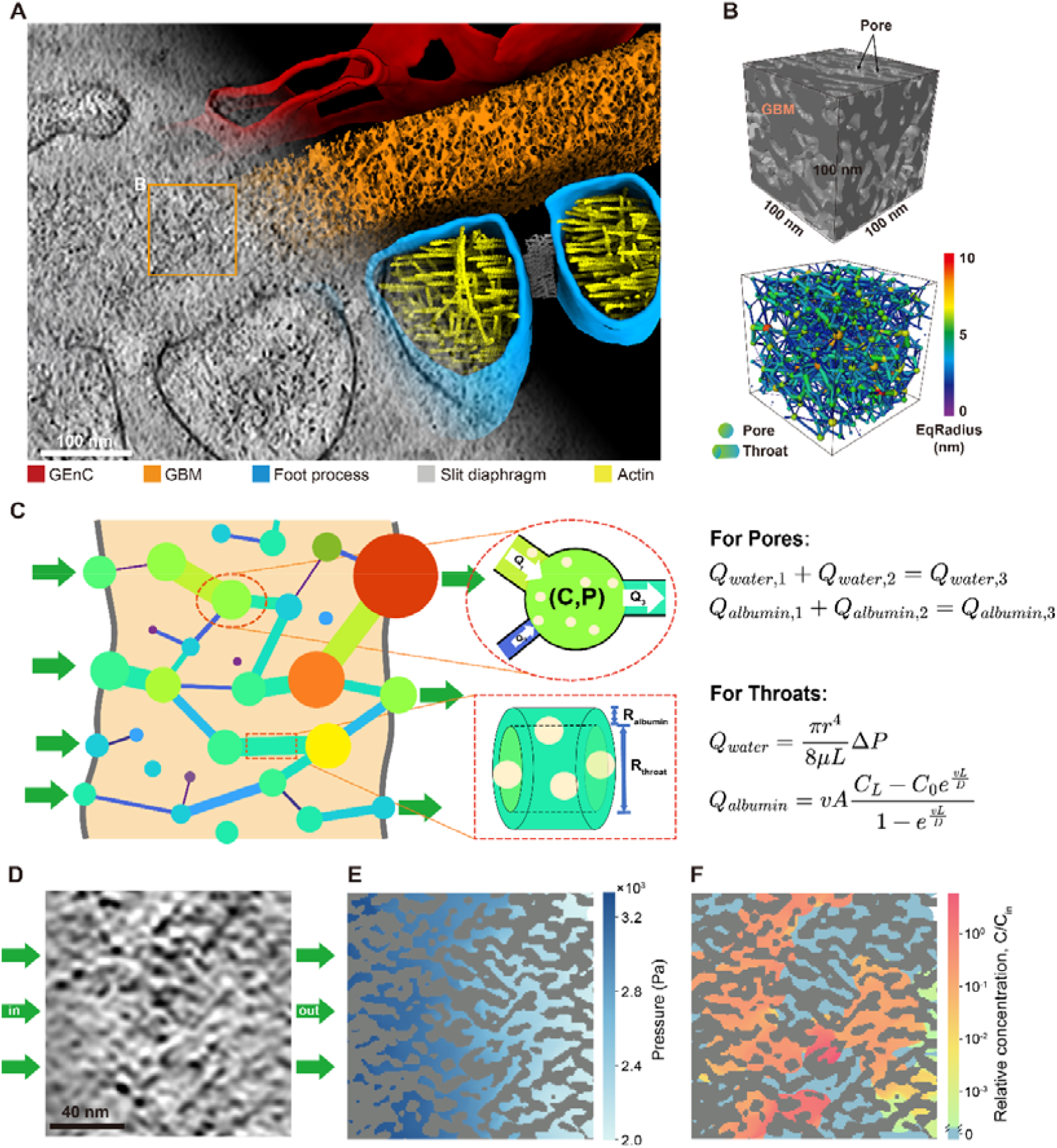
Structural basis and transport modeling of albumin sieving in the glomerular basement membrane. (A) Cryo-ET reconstruction of the glomerular filtration barrier, shown as a transition from the raw tomographic slice to the corresponding 3D segmentation. The reconstruction delineates the spatial relationship between endothelial cells (GEnCs), the GBM, and podocyte foot processes. The orange box marks a representative region within the GBM analyzed in (B). (B) 3D segmentation and pore-network representation of the GBM. The upper panel shows the segmented volume, with the fibrillar matrix shown in dark gray and inter-fibrillar void space rendered as transparent regions. The lower panel displays the corresponding pore-network model, in which color-coded spheres represent pores, and connecting lines represent throats connecting adjacent pores. Equivalent radius is color-coded as indicated. (C) Schematic of the pore-throat transport model used to calculate GBM albumin sieving. Pores are represented as nodes and throats as connecting edges with node and edge colors indicating equivalent radius as in (B). For each pore, steady-state solvent volume flow rate and albumin volume flow rate are conserved. P denotes steady-state nodal pressure and C denotes the steady-state equivalent albumin concentration at pore nodes. For each throat, solvent volume flow rate *Q*_*water*_ is calculated from Poiseuille flow, and albumin volume flow rate*Q*_*albumin*_ is calculated from the steady one-dimensional advection-diffusion solution. Here, *r* is throat radius, *L* is throat length, μ is water viscosity. *v* is axial albumin velocity, *A* is albumin accessible area and *D* is the effective albumin diffusivity of a throat. (D) A representative cross-section of tomogram collected in GBM region. Electron-dense fibrillar matrix appears gray, whereas inter-fibrillar pore space appears bright. Green arrows indicate flow direction. (E) Calculated pressure distribution in the same region shown in (B) and (D). Gray regions indicate the segmented fibrillar matrix. Colored void regions indicate the steady-state pressure field calculated from the pore-throat solvent-flow model. (F) Calculated relative albumin concentration distribution in the same region shown in (B) and (D). Gray regions indicate the fibrillar matrix, and blue regions indicate void space that is inaccessible to albumin under the radius threshold. Colored void regions show the calculated relative albumin concentration, normalized to the inlet albumin concentration *C*_*in*_. Scale bars: 100 nm in A; 40 nm in D.

Using these pore-throat maps as the structural input, we built a structure-resolved transport model (see Methods). Unlike idealized models based on uniform cylindrical pores ^35,36^ or random fiber arrays ^37,38^, this approach preserves nanoscale structural parameters measured from *in situ* tomograms, including throat radius, throat length, and network connectivity, to calculate solvent flow and albumin transport through the reconstructed GBM geometry (Fig. 4C).

The same pore-throat geometry places solvent and albumin in distinct transport regimes. For solvent, the GBM network behaves as a well-connected hydraulic medium: solving flow through the reconstructed network produced a smooth, monotonic pressure drop from the capillary side toward Bowman’s space (Fig. 4E). Albumin transport, by contrast, lies in a sterically restricted regime. Quantitative analysis of cryo-ET reconstructions showed that approximately 80% of GBM throat radii are smaller than the albumin hydrodynamic radius, eliminating these connections from the albumin-accessible network. The remaining accessible space was therefore reduced to sparse connected paths composed entirely of albumin-permissive throats (Fig. 4F).

We next solved albumin advection-diffusion on this albumin-accessible subnetwork under the local flow field. Unlike solvent pressure, albumin concentration did not vary smoothly across the GBM. Even on a logarithmic scale, the concentration profile showed sharp drops across narrow throats. These bottlenecks imposed high local resistance under flow, further suppressing flux along the already sparse albumin-accessible paths. Direct calculations on the finite reconstructed networks yielded a mean sieving coefficient of approximately 0.003 (Extended Data Fig. 7), corresponding to more than 99.7% reduction relative to the inlet concentration.

### GBM pore-network remodeling weakens albumin retention in Alport syndrome

We next asked whether the proposed GBM sieving mechanism could explain filtration failure in disease. We examined the GFB in 8-week-old *Col4a3*^−/−^ mice at an early proteinuric stage preceding severe glomerular scarring. The overall GFB architecture remained recognizable at this stage (Fig. 5A–D; Extended Data Fig. 8A–E; Supplymentary Movie 5 and 6). Across 426 tomograms, endothelial fenestral dimensions and slit-diaphragm width and architecture remained broadly comparable to those in healthy glomeruli at the current resolution. The dominant detectable structural changes were instead concentrated in the GBM (Extended Data Fig. 9).

At the mesoscale, the lamina densa showed pronounced thickness heterogeneity, including substantial local thickening (Fig. 5C–D). If the thickened regions retained healthy matrix architecture, the greater crossing distance would reduce the probability of continuous albumin-accessible routes and add transport bottlenecks along any remaining route. This expectation constrasts with the albumin leakage observed in Alport syndrome.

**Fig. 5.**
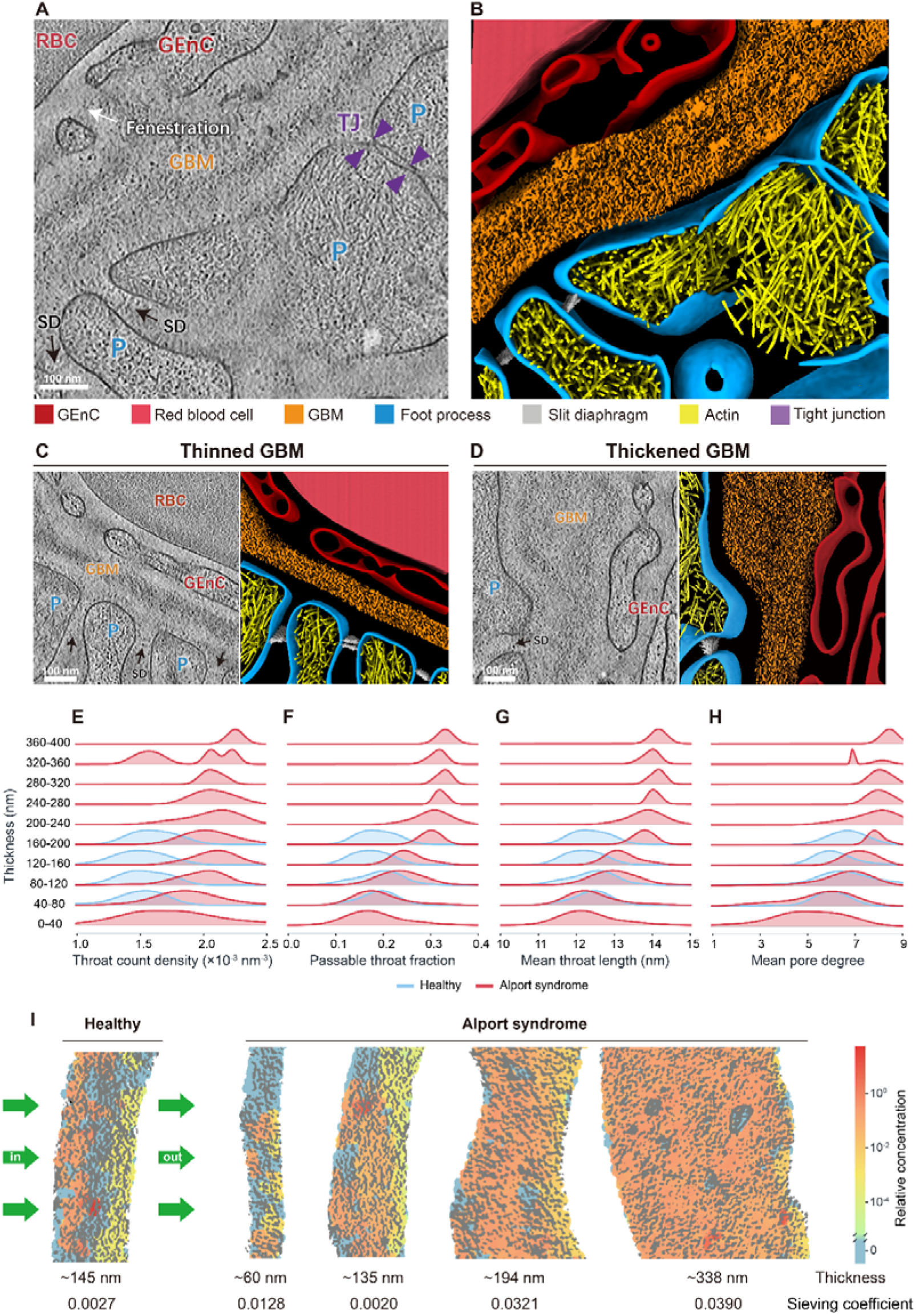
Glomerular basement membrane remodeling increases albumin permeability in Alport syndrome. (A–B) Representative tomographic slice of the glomerular filtration barrier collected from Alport syndrome mice (A) and the corresponding 3D segmentation of subcellular structures as indicated (B).(C–D) Representative abnormal GBM regions in Alport syndrome mice, showing focal GBM thinning (C) and thickening (D). (E–H) Thickness-stratified distributions of pore-throat structural descriptors from GBM subsamples. Ridgeline plots show throat count density (E), albumin-passable throat fraction (F), mean throat length (G), and mean pore degree (H) across GBM thickness bins. Healthy subsamples are shown in blue and Alport syndrome subsamples in red. Albumin-passable throats were defined using the hydrated albumin radius threshold of 3.55 nm. Descriptor distributions were calculated from 177 Healthy subsamples and 283 Alport syndrome subsamples. (I) Representative calculated relative albumin concentration maps in one Healthy GBM network and four Alport syndrome GBM networks spanning increasing GBM thickness. Approximate GBM thicknesses were 145 nm for the Healthy sample and 60, 135, 194, and 338 nm for the Alport syndrome samples. The corresponding calculated albumin sieving coefficients were 0.0027 for the Healthy sample and 0.0128, 0.0020, 0.0321, and 0.0390 for the Alport syndrome samples. Gray regions indicate GBM matrix, blue regions indicate albumin-inaccessible void space, and colored albumin-accessible regions show relative albumin concentration normalized to inlet albumin concentration. Green arrows indicate flow direction. Scale bars: 100 nm in A, B, C and D.

At the nanoscale, cryo-ET analysis resolved this paradox by revealing thickness-dependent remodeling of GBM topology. Thicker Alport syndrome regions exhibited greater throat count density, a larger fraction of albumin-accessible throats, longer throats and higher pore degree (Fig. 5E–H), indicating a more open and connected void network. Transport modeling further showed that as thickness increases, the bottleneck pattern in albumin concentration weakened markedly. In the thickest samples, albumin became broadly and uniformly distributed, showing passage along numerous connected routes (Fig. 5I). Correspondingly, these samples yielded the highest sieving coefficients, providing a mechanistic explanation for albumin leakage in Alport syndrome. In turn, the Alport syndrome model serves as a perturbation test that further supports the topology-driven mechanism of albumin sieving by the GBM.

### Ensemble networks connect local architecture to GBM-scale sieving

Cryo-ET resolves native structures locally, whereas tissue physiology emerges from their collective behavior across much larger scales. Glomerular filtration presents this local-to-global problem directly: overall albumin selectivity should reflect structural heterogeneity throughout the GBM (Fig. 6A–B). To bridge these scales, we developed a statistical ensemble approach. We first characterized the distributions of microstructural descriptors, including throat radius, throat length, throat density and pore degree, and their dependence on local GBM thickness in the cryo-ET volumes (Fig. 6C). We then combined these nanoscale statistics with the measured mesoscale distribution of GBM thickness to generate ensembles of GBM networks (Extended Data Fig. 6, Extended Data Fig. 10 and Extended Data Fig. 11; see Methods). The generated networks reproduced the measured structural statistics without fitting to physiological sieving, enabling an independent prediction of overall albumin passage.

**Fig. 6.**
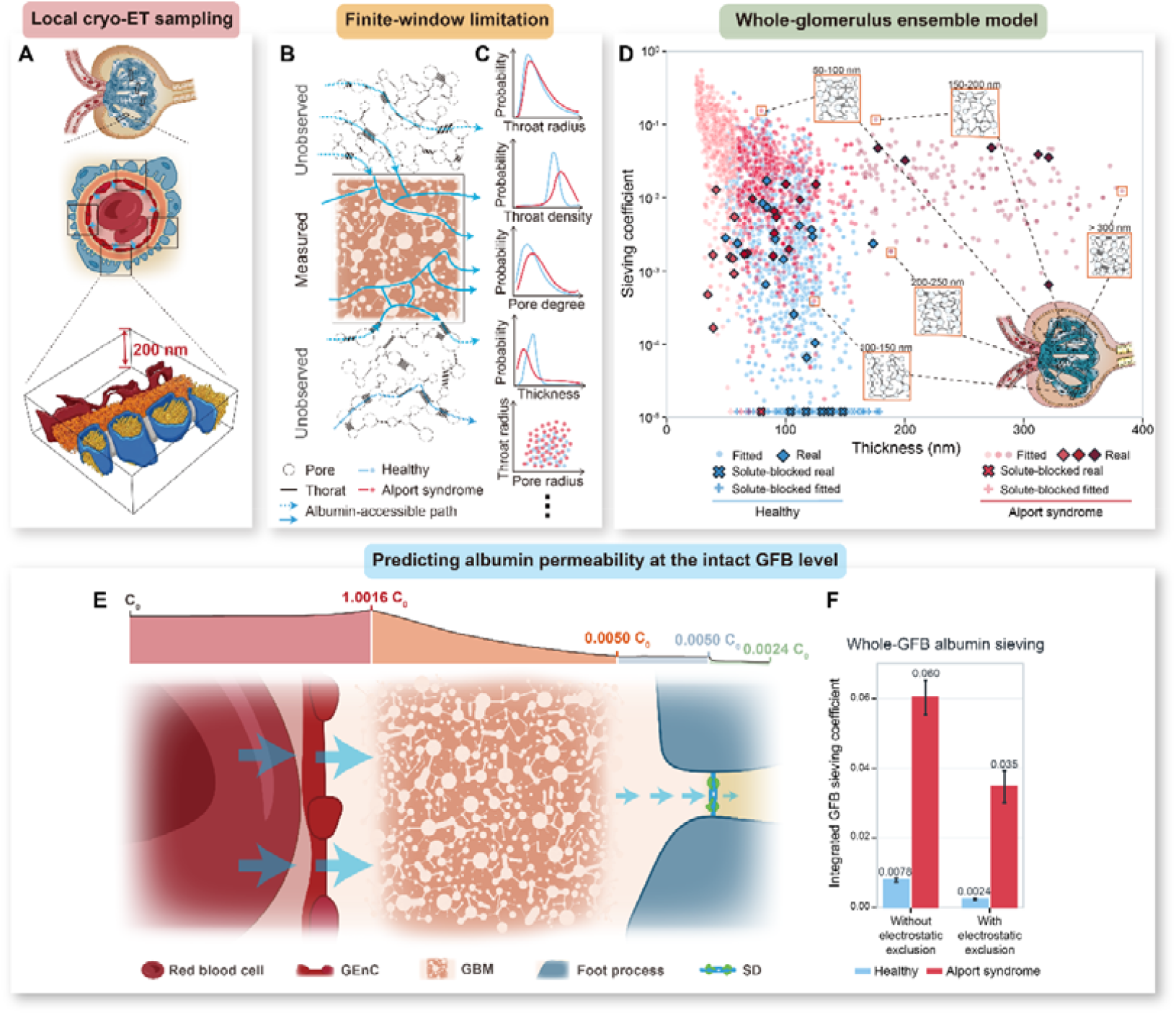
Fitted glomerular basement membrane network model and whole-glomerulus albumin sieving prediction. (A) Schematic of the local cryo-ET sampling workflow. Tilt-series data were acquired at multiple positions across glomerular capillary regions, yielding tomographic reconstructions spanning approximately 200 nm along the lamella thickness. (B) Each acquired tomographic dataset preserves a transverse filtration pathway from the capillary lumen to Bowman’s space, but some regions remain incompletely sampled owing to lamella thickness or acquisition-window constraints, including the pore-throat network indicated by dashed lines. Solid blue lines denote observed albumin-accessible pathways; dashed blue lines denote albumin-accessible pathways outside the sampled volume. (C) Probability distributions of representative microstructural descriptors extracted from Healthy and Alport syndrome GBM samples, including throat radius, throat density, pore degree, and corrected GBM thickness. A representative descriptor–descriptor correlation between pore and throat radii is shown below. Blue and red indicate Healthy and Alport syndrome samples, respectively. (D) Thickness-dependent albumin sieving coefficients predicted by the GBM transport model. Healthy networks are shown in blue, and Alport syndrome networks are shown in red. Red shades from light to dark correspond to fitted GBM thickness intervals of less than 60 nm, 60 to 160 nm and greater than 160 nm. Diamonds indicate values calculated directly from reconstructed real GBM networks, whereas circles indicate fitted synthetic networks. Each circle corresponds to a network fitted at a specific thickness and represents a distinct region of the GBM. The orange box shows fitted results for five regions with different thicknesses. Cross markers at the lower axis floor indicate solute-blocked networks with sieving coefficient lower than 10^-5^ or zero sieving. (E) Whole-barrier prediction of albumin sieving across the intact healthy GFB. Albumin transport was modeled as four sequential steps from capillary plasma to Bowman’s space: two short bulk convection-diffusion regions, the GBM, and the slit diaphragm (SD). Concentrations were normalized to plasma albumin concentration *C*_0_. The four-layer calculation yielded concentration ratios of 1.0016 for the plasma-to-GBM boundary layer, 0.005 for the GBM layer, 1.0008 for the GBM-to-slit diaphragm cleft, and 0.48 for the slit diaphragm lattice protein layer. The combined model predicted *C*_*Bowan*_=0.0024*C*_0_, corresponding to an intact GFB albumin sieving coefficient of 0.0024. (F) Predicted integrated albumin sieving coefficients of the intact GFB in Healthy and Alport syndrome ensembles, calculated without or with electrostatic exclusion. Whole-GFB values were obtained by multiplying the integrated GBM sieving coefficients by the fixed transfer factors of the plasma-to-GBM region, GBM-to-slit diaphragm cleft, and slit diaphragm. Without electrostatic exclusion, the coefficients were 0.0078±0.0006 for Healthy ensembles and 0.060±0.005 for Alport syndrome ensembles; with electrostatic exclusion, they were 0.0024±0.0003 and 0.035±0.005, respectively. Bars show the integrated GFB albumin sieving coefficients; error bars indicate ± standard deviation across 200 thickness-stratified bootstrap estimates.

Transport modeling quantified albumin sieving for each generated network in the Healthy and Alport syndrome ensembles (Fig. 6D). Flux-weighted averaging across each ensemble yielded integrated sieving coefficients of 0.016 ± 0.001 for Healthy GBM and 0.13 ± 0.01 for Alport syndrome GBM, an approximately eightfold increase in disease (Extended Data Fig. 12). The higher coefficient arose from leakage through both thickened and thinned GBM regions. Leakage in the thin regime was underestimated by direct calculations on cryo-ET samples, which are finite slices that can truncate laterally extended routes and make continuous paths appear disconnected. The generated networks reduced this finite-window bias and showed that shorter crossing distances increase the likelihood of continuous albumin-accessible routes. Sensitivity analysis further identified throat radius as the most consistent structural driver of increased Alport syndrome sieving, with throat density also contributing (Extended Data Fig. 13; Extended Data Table 2A).

Considering albumin and the GBM are both negatively charged at physiological pH ^39,40^, we next studied the contribution of electrostatic repulsion. We incorporated this effect into the pore–throat model using an effective-radius approximation, increasing the albumin exclusion radius by one Bjerrum length in water, from 3.55 to 4.25 nm ^41^. This correction reduced the integrated sieving coefficient from 0.016 to 0.005 in Healthy GBM and from 0.13 to 0.072 in Alport syndrome GBM (Extended Data Fig. 14). The fraction of networks permitting albumin passage also decreased from 74.6% to 28.9% in the Healthy ensemble and from 99.6% to 85.0% in the Alport syndrome ensemble (Extended Data Fig. 14). Thus, electrostatic repulsion strengthened albumin restriction without erasing the topology-driven difference between Healthy and Alport syndrome GBM.

### Serial integration of GFB layers predicts physiological albumin sieving

Direct validation of the layer-specific predictions would require isolating each GFB layer without disrupting its native architecture. We therefore used a glomerular albumin sieving coefficient of 0.0023, previously measured in isolated perfused mouse kidneys, as an independent benchmark ^42^. To generate a whole-barrier prediction, we coupled the structure-derived transport estimates for each layer in their physiological sequence from capillary plasma to Bowman’s space (Fig. 6E; see Methods). Endothelial fenestrae were treated as nonrestrictive to albumin based on their measured dimensions. GBM sieving was obtained from the generated network ensemble and slit-diaphragm sieving from the measured distribution of slit widths, while the intervening fluid regions were treated as bulk transport compartments without geometric sieving.

The serial model enabled a direct comparison of albumin restriction across the GFB. The predicted albumin concentration changed little across the bulk fluid regions, dropped sharply across the GBM, and decreased more modestly across the slit diaphragm (Fig. 6E). This further confirms the major role of GBM in albumin sieving. At the whole-barrier level, the model predicted a sieving coefficient of 0.0078 under steric exclusion alone and 0.0024 after incorporating electrostatic exclusion in the GBM (Fig. 6F). Both values fell within the same order of magnitude as the independent benchmark of 0.0023, with the latter even closely matching its value. This agreement demonstrates that locally measured architecture can be integrated across scales to predict physiology at the tissue level. It also shows that steric exclusion imposed by GFB architecture is the principal determinant of albumin sieving and electrostatic repulsion strengthens this restriction.

## Discussion

By combining cryo-ET of vitrified glomeruli with structure-resolved transport modeling, we extend structure-function analysis from molecular assemblies to an intact tissue barrier. Two complementary steps make this possible. First, geometry supplies the physical basis: experimental reconstructions specify the local pathways available to solvent and albumin, while established transport laws and physiological gradients govern the resulting fluxes. Second, a generative rule bridges spatial scales: thickness-dependent network statistics extracted from local cryo-ET volumes, combined with the mesoscale distribution of GBM thickness, produce ensembles representing tissue-scale heterogeneity. Without calibration to the physiological sieving coefficient, the resulting whole-barrier model recovers the measured scale of albumin retention. Native tissue architecture can therefore serve as a quantitative basis for predicting filtration function.

This structure-to-physiology framework reveals that albumin retention arises primarily from the GBM. The underlying mechanism lies in the topology of its inter-fibrillar void space. Water traverses a broadly connected network, whereas steric exclusion renders narrow throats inaccessible to albumin and confines it to sparse paths punctuated by transport bottlenecks. This size selectivity is a collective, percolation-like property of the void network rather than a function of a single characteristic pore dimension. This topological information is absent from physiological models that reduce the GBM to an effective pore size or fibre density^35–38^. Indeed, substituting the cryo-ET-measured mean pore radius of 4.6 nm and matrix volume fraction of 0.55 into such models predicts albumin sieving on the order of 10^−6^, far below the physiological scale. Detailed native three-dimensional geometry is therefore essential for predicting GBM filtration from structure.

Alport syndrome provides a disease test of this topology-dependent mechanism. At the early proteinuric stage examined here, structural changes were concentrated in the GBM, while endothelial fenestrae and slit-diaphragm architecture remained broadly preserved. The molecular defect lies in a failed developmental switch: the embryonic α1α1α2 collagen-IV network is normally replaced by the more highly crosslinked α3α4α5 network, which confers greater proteolytic resistance and mechanical stability ^43,44^. In *Col4a3*^*−/−*^ mice, α1α1α2 instead persists as an incomplete substitute ^45,46^. The resulting instability manifests in increased GBM thickness heterogeneity and remodeling of its internal void network. Both thinned and thickened regions promote albumin passage, but through distinct topological mechanisms: thinning shortens the crossing path, whereas thickening is associated with increased network connectivity. At later stages, slit-diaphragm displacement, ectopic tight-junction formation and fibrillar collagen deposition accompany the loss of recognizable GFB organization ^47^. Together, these observations place GBM topological disruption early in the broader breakdown of filtration architecture, although static reconstructions cannot establish the temporal sequence or causality of these changes.

Unlike the GBM relying on irregular network topology for permselectivity, the two cellular layers contain ordered molecular assemblies suited to distinct functions. At endothelial fenestrae, a putative EHD-family rim complex is formed to stabilize openings at a functional size, large enough to permit plasma flow but small enough to exclude circulating cells. At the slit diaphragm, a flexible Nephrin-Neph1 lattice bridges adjacent foot processes but provides insufficient resistance to function as a stringent albumin sieve. This does not diminish the biological importance of the slit diaphragm; instead, it points to a primary role in extracellular adhesion and podocyte organization. Pathogenic *NPHS1* variants map near predicted Nephrin-Nephrin interfaces, suggesting that impaired lattice assembly or stability may contribute to disease. Cytoplasmic densities closely associated with actin filaments beneath the slit diaphragm (Extended Data Fig. 15), together with established Nephrin-Nck-N-WASP signalling pathway ^48,49^, further suggest that the extracellular lattice is coupled to the podocyte cytoskeleton. Disruption of either extracellular lattice integrity or its cytoskeletal coupling could therefore destabilize foot-process architecture and cause proteinuria without requiring the slit diaphragm to act as a stringent molecular sieve.

A complete molecular reconstruction of a tissue is not feasible and may not be necessary to predict physiology. What is required is a statistically faithful description of the structures governing the process. Here, cryo-ET-derived void-network statistics and the GBM thickness distribution provide such a description at the tissue scale. More generally, local high-resolution structures can be integrated with coarser measurements of tissue heterogeneity to generate representative ensembles. Physical modelling can then translate these ensembles into function. The present analysis simplifies the in vivo mechanical environment and spatially heterogeneous electrostatic interactions. Larger reconstructions and systematic size- and charge-dependent tracer measurements can test and refine these approximations. This local-to-ensemble strategy could extend cryo-ET from resolving molecular assemblies to predicting collective physiology in other biological barriers, extracellular matrices and multicellular interfaces.

## Acknowledgments

We thank the Cryo-EM Platform of Peking University and Changping Laboratory, the National Centre for Protein Sciences at Peking University, and the National Multimode Trans-scale Biomedical Imaging Center for technical assistance.

This work was funded by the National Key Research and Development Program of China (#2024YFA1802800 to Q.G.), the Beijing Natural Science Foundation (JQ24031 to Q.G.), and the National Natural Science Foundation of China (#32371191 to Q.G.). Q.G. is supported by Changping Laboratory and the SLS-Qidong Innovation Fund. Q.G. and M.H. acknowledge supports from the Peking-Tsinghua Center for Life Sciences.

## Declaration of interests

The authors declare no competing interests.

## Data and code availability

All data supporting the findings of this study are available within the paper and its Supplementary Materials. The subtomogram averaging maps have been deposited to the Electron Microscopy Data Bank (EMDB) under accession codes EMD-82753 for the slit diaphragm and EMD-82754 for the fenestral rim complex. Representative tomograms of the glomerular filtration barrier in healthy and Alport syndrome mice have been deposited to EMDB under accession codes EMD-82755 and EMD-82756, respectively. The source code for numerical simulations is available on GitHub (https://github.com/zzz721/GFB-albumin-sieving-model.git).

## Methods

### Experimental animal ethics and management

All experimental procedures involving animals received explicit approval from the Institutional Animal Care and Use Committee (IACUC) of Peking University First Hospital, operating under the designated Animal Ethics Committee Approval Number: 2025181. Adherence to the stringent ethical guidelines outlined in the National Institutes of Health (NIH) Guide for the Care and Use of Laboratory Animals was strictly maintained throughout the study. Furthermore, all aspects of animal experimentation, data reporting, and welfare considerations were meticulously conducted and documented in full compliance with the ARRIVE guidelines (Animal Research: Reporting of In Vivo Experiments).

### Animal model and housing conditions

The study utilized *Col4a3* knockout (*Col4a3*^*-/-*^) mice, which were generated on a C57BL/6J genetic background (C57BL6/J-*Col4a3*^tm1Dec^) and generously provided by Sanofi-Genzyme. The C57BL/6J strain is a widely recognized and well-characterized laboratory mouse strain frequently employed in metabolic, cardiovascular, and renal research due to its robust genetic background and established disease models. The *Col4a3*^*-/-*^ mouse model is particularly relevant for studying Alport syndrome, a progressive genetic kidney disorder characterized by defects in type IV collagen, which is a major component of the glomerular basement membrane. These mice develop progressive glomerulopathy and proteinuria, mimicking the human disease phenotype.

All *Col4a3*^*-/-*^ mice were housed in a specific pathogen-free (SPF) environment within a dedicated laboratory animal facility. The housing conditions included a controlled 12-hour light/dark cycle and a stable room temperature of 25□, designed to minimize environmental stress and maintain physiological homeostasis in the animals. Consistent with best animal care practices, both food and water were provided ad libitum, allowing unrestricted access to nutrition and hydration.

### Sample collection

To facilitate comprehensive longitudinal analyses of disease progression and treatment effects, biological samples were systematically collected at predetermined time points: 8 and 18 weeks of age. This staggered collection strategy enables the capture of different stages of disease development and potential therapeutic responses. Urinary samples, critical for assessing renal function and proteinuria, were collected using metabolic cages. Metabolic cages allow for the precise and uncontaminated collection of urine over a defined period (24 hours in our study), enabling quantitative measurements of urinary analytes such as albumin and creatinine. This method ensures accurate assessment of kidney function, as exemplified by urinary albumin-to-creatinine ratio (uACR) measurements, which are highly sensitive indicators of glomerular injury.

Kidney tissues were also harvested at these time points for subsequent histological and ultrastructural examinations, providing direct insights into the structural and cellular changes associated with the disease. Glomeruli were isolated via mechanical sieving, a well-established method enabling high-purity enrichment ^20^. Briefly, mice were euthanized by cervical dislocation, and kidneys were immediately excised and immersed in ice-cold PBS. After removal of the renal capsule and perirenal fat, tissues were gently minced and sequentially filtered through a cascade of sterile cell strainers (100 μm, 70 μm, 40 μm) to enrich for glomerular fragments. The glomerular fraction retained on the 40 μm filter was collected, washed twice in cold PBS, and resuspended in fresh buffer for downstream applications.

### Urine albumin to creatinine ratio measurement

The urinary albumin concentration was quantitatively assessed utilizing a Mouse Albumin Enzyme-linked Immunosorbent Assay (ELISA) Kit (Bethly, E99-134). This methodology relies on a highly sensitive immunoassay principle, enabling precise detection of albumin in biological samples. Concurrently, urinary creatinine levels were determined using a QuantiChrom™ Creatinine Assay Kit (DICT-500, BioAssay Systems). The determination of creatinine serves as an essential internal control for normalizing urine output variations. Subsequent to individual measurements, the uACR values were computed and are presented in units of micrograms of albumin per milligram of creatinine (µg albumin/mg creatinine). This standardized ratio provides a robust indicator of glomerular protein filtration, independent of urine dilution or concentration variability.

### Immunofluorescence staining

For immunofluorescence analysis, glomerular sections underwent a series of preparatory steps to optimize antibody binding and signal detection. Initially, the sections were dewaxed and rehydrated through a graded ethanol series, followed by three washes with PBS to remove residual fixatives. To minimize non-specific antibody binding, sections were then subjected to a blocking step using 1% BSA in PBS for 1 hour at 25□. Primary antibodies targeting key podocyte proteins were applied: rabbit anti-mouse podocin (dilution 1:5000, Abcam, ab50339) and mouse anti-mouse synaptopodin (dilution 1:100, Progen, 61094). Incubation with these primary antibodies was carried out overnight at 4□ to facilitate specific antigen-antibody interactions. Following extensive washing to remove unbound primary antibodies, sections were incubated for 1 hour at 25□ with highly cross-adsorbed secondary antibodies conjugated to Alexa Fluor™ 594: donkey anti-rabbit IgG (H+L) (dilution 1:500, Invitrogen, A21207) and goat anti-mouse IgG (H+L) (dilution 1:500, Invitrogen, A-11005). Alexa Fluor™ 594 is a spectrally stable fluorophore suitable for multiplex imaging. The stained sections were then mounted onto coverslips using a mounting medium containing DAPI (Zhongshan Golden Bridge Biotechnology, Beijing, China), which serves as a nuclear counterstain. Image acquisition was performed using a confocal microscope (Zeiss 780, Germany), allowing for high-resolution imaging of fluorescent signals within the glomeruli.

### Transmission electron microscopy

For ultrastructural examination by transmission electron microscopy (TEM), tissue samples were processed with meticulous attention to preserve details. The initial fixation involved immersion in 2.5% glutaraldehyde at 4□ overnight, a strong cross-linking agent that stabilizes protein structures. Following primary fixation, samples were washed three times in 0.1 M cacodylate buffer to remove excess glutaraldehyde. Post-fixation was performed using 1% osmium tetroxide for 2 hours, which enhances membrane contrast and further stabilizes lipids. Subsequently, samples underwent dehydration through a series of increasing acetone concentrations (30%, 50%, 70%, 80%, 90%, and 100% acetone solution) to gradually remove water and prepare the tissue for resin embedding. Ultra-thin sections, precisely 50 nm in thickness, were then cut using an ultrathin microtome (Leica EM UC7, Wetzlar, Germany). To enhance contrast for electron visualization, these sections were stained with 2% uranyl acetate for 10 minutes, followed by lead citrate for 5 minutes. The final observation and imaging of the ultrastructural features were conducted using a TEM (JEM-1400, JEOL Ltd., Tokyo, Japan), providing detailed insights into the nanometer-scale morphology of the glomerulus.

### High pressure freezing

Glomerular samples were prepared for cryo-electron tomography (cryo-ET) using an optimized vitrification protocol. Briefly, EM grids (Quantifoil, Cu, 200 mesh, R2/1) were subjected to glow discharge treatment to enhance hydrophilicity. Under stereomicroscopic observation, individual glomeruli were carefully transferred from the dish to the grid center using self-locking tweezers. To optimize sample preparation, excess liquid was gently absorbed from the dorsal side of the grid using filter paper, followed by the application of 4 µL of cryoprotectant solution (5% sucrose and 20% dextran in PBS). The samples were then incubated for 30 s to ensure proper cryoprotectant penetration, after which residual liquid was carefully removed from the dorsal side. Then, the grid was quickly placed at the center of a 6 mm aluminum carrier (200 µm in depth), which had been pre-wetted with 2-methylpentane (Sigma). After the carrier was filled with 2-methylpentane, a 6 mm sapphire disc was promptly placed on top. Freezing was initiated immediately using a Leica EM ICE high-pressure freezing system after the sandwich carrier was assembled, and only samples with intact sapphire discs were selected for further processing. For obtaining non-embedded grids, the sandwich carrier was rapidly transferred from liquid nitrogen into a cryogenic mixture of propane and ethane maintained at −170□ for 2 min using the temperature-controlled chamber of the Leica EM-GP plunger. Following cryo-immobilization, the carrier was carefully disassembled under controlled conditions.

### Serial lift-out and FIB thinning

Cryo-focused ion beam (cryo-FIB) lift-out and lamella preparation were carried out using an Aquilos 2 cryo-FIB system (Thermo Fisher Scientific), following protocols established in our prior work ^21^. Briefly, both the sample-containing grid and slotted mesh grid (G100/400) were loaded onto the shuttle. To mitigate sample charging and ensure sufficient surface conductivity, the grid was subjected to gas-assisted platinum deposition for 30 s followed by sputter coating at 30 mA for 15 s. Initial milling was conducted using a Regular Cross Section Pattern with a 30 kV Ga^2+^ ion beam at 3 nA current. The stage was then reoriented to align the tissue surface perpendicular to the ion beam, enabling precise planarization of the region of interest at a reduced beam current of 1 nA. For lift-out, an EasyLift micromanipulator needle affixed with a copper support chunk was inserted into the chamber. The bottom face of the copper chunk was polished with a 5 nA ion beam, brought into contact with the pre-milled sample surface, and bonded via ion-beam-assisted platinum redeposition at 1 nA. The targeted lamella was then excised using a Line milling pattern, transferred to the support grid, and securely anchored through additional platinum redeposition. This lift-out cycle was iteratively repeated to fully dissect the tissue block.

Following extraction, lamellae were further thinned using a multi-step milling protocol. Coarse milling began at 1 nA and was progressively refined by reducing the ion beam current as the target thickness approached, thereby minimizing ion-induced damage. Final polishing was performed at 50–30 pA to yield electron-transparent lamellae with a thickness under 200 nm. The process concluded with a brief sputter coating at 7 mA for 10 s to enhance surface stability during subsequent cryo-ET imaging.

### Serial cryo-SEM volume imaging

Glomerular tissue blocks measuring approximately 40 × 40 × 15 μm^3^ were extracted via cryo-focused ion beam lift-out (cryo-FIB lift-out) and transferred onto standard TEM support grids. Serial FIB/SEM volume imaging was performed on a Thermo Fisher Scientific Helios 5 Hydra CX dual-beam system using Auto Slice & View 5 software (Thermo Fisher Scientific).

Grids were mounted on a 35° pre-tilted holder and sequentially coated with conductive and protective layers. First, platinum was deposited by magnetron sputtering at 12 kV and 0.25 microamperes for 60 s. Next, organometallic platinum was applied via a gas injection system for 240 s. Finally, a second platinum sputtering layer was deposited at 12 kV and 0.25 microamperes for 300 s.

For image acquisition, the stage was tilted to 35° with a working distance of 4.0 mm to orient the grid surface perpendicular to the electron beam, resulting in a 52° FIB milling angle relative to the sample surface. A fiducial trench was milled adjacent to the region of interest to enable drift correction during long-duration acquisitions. Ion milling was carried out using a 30 kV Ar□ beam at a nominal current of 0.2 nA, with a slice thickness of 20 nm.

SEM imaging was conducted in immersion mode using a through-lens detector (TLD) operating in high-gain diode mode. Imaging parameters were tailored for each sample type to optimize for contrast and stability: the Alport syndrome sample was imaged at 1.2 kV and 13 pA, with a 100 ns dwell time, 90-line integration, a pixel size of 6 × 6 nm, and a resolution of 3500 × 2700 pixels; the Healthy sample was imaged at 1.0 kV and 13 pA, with a 50 ns dwell time, 110-line integration, a pixel size of 5 × 5 nm, and a resolution of 4200 × 4000 pixels. During acquisition, auto-focus and auto-stigmatism routines were executed after every fifth image.

### Volume imaging processing and segmentation

Acquired 2D serial section images were first imported into Amira 2024 (Thermo Fisher Scientific) for automated alignment. To reduce noise while preserving edge fidelity, images were smoothed using the Anisotropic Diffusion filter. Residual periodic stripe artifacts (horizontal and vertical) were subsequently removed via the FFT Filter module. The processed 2D images were exported as TIFF stacks.

To enhance local contrast, images were further processed in Fiji using the Enhance Local Contrast (CLAHE) function, with parameters optimized empirically for each dataset. The contrast-enhanced stacks were re-imported into Amira for refined alignment. This iterative processing alignment cycle was repeated 2–3 times to achieve optimal registration and artifact suppression.

3D segmentation of the glomerular ultrastructure was performed in Amira 2024 using AI-assisted segmentation based on a deep learning model. A ResNet18 model was employed as the backbone for the segmentation network, which demonstrated high accuracy in identifying endothelial fenestration. The initial AI-generated labels were manually modified, ensuring accuracy of the final 3D reconstructions.

### Cryo-ET data acquisition

Cryo-ET data for Healthy and Alport syndrome glomerulus were acquired on a 300□kV Titan Krios G4 microscope (Thermo Fisher Scientific) equipped with an X-FEG electron source and a Falcon 4i direct electron detector (Thermo Fisher Scientific). Tilt series were recorded using Tomography 5 software (Thermo Fisher Scientific) with a dose-symmetric tilt scheme ^50^ over an angular range of −40° to +60° (relative to the pre-tilt angle), using 2° increments. An energy-selecting slit with a 10 eV full width was employed to enhance contrast by filtering inelastically scattered electrons. Imaging was performed in electron-counting mode at a calibrated pixel size of 2.37□Å, with defocus values maintained between −2.0 and −6.0□μm. The total dose for each tilt series was maintained at 120 e^−^/Å^2^ and dose rate on the detector was targeted to be ~8–9 e^-^ /pix/s.

### Tomogram processing pipeline

Individual projection images were motion-corrected using MotionCor2 ^51^. Low-quality projection images were excluded based on visual inspection. Tilt series from Healthy samples 001 to 462 were aligned using the patch-tracking algorithm implemented in IMOD 4.11.3 ^52^. Subsequent tilt series, comprising Healthy samples 463 to 549 and Alport syndrome samples 001 to 426, were processed with patch-tracking alignment in AreTomo2 ^53^. Contrast transfer function (CTF) parameters were estimated, and tomograms were reconstructed by weighted back-projection using Warp 1.0.9 ^54^.

To improve signal-to-noise ratio, tomograms were denoised with IsoNet v0.2 ^55^ at an effective voxel size of 9.48□Å/pixel with 4× binning. Denoised, 4× binned tomograms served as input for membrane segmentation via MemBrain v2 ^56^. The resulting segmentation masks were imported into Amira 2020.3.1 (Thermo Fisher Scientific), where any missing or ambiguous membrane regions were manually corrected and completed.

### Subtomogram averaging of the fenestral rim complex

Initial particle picking was performed manually on 10 tomograms containing endothelial cells. Helical segments were oversampled using the *relion_helix_toolbox* in RELION/3.1, applying a helical rise of 38□Å per repeat, yielding 1,532 particle segments. These particles, binned by 2 at a box size of 96 pixels, were extracted using Warp 1.0.9 and averaged in RELION, using an initial reference template modeled as a rectangular slab matching the thickness of the plasma membrane. 3D classification revealed a distinct class corresponding to the fenestral rim complex with clear helical periodicity, comprising 10 dimeric units.

This reference was then used to guide alignment in an expanded dataset, including 16 Healthy and 59 Alport syndrome tomograms. All newly picked particles were merged with the initial set for joint refinement. During 3D classification, a soft mask was applied exclusively to the fenestral rim complex region to exclude adjacent membrane densities. Particles assigned to the dominant class, accounting for 75% of the total, were re-extracted at full pixel size with a box size of 120 pixels for high-resolution alignment.

To improve angular accuracy, aligned particles were imported into ChimeraX-1.10.1 and refined using ArtiaX plugin ^57,58^, which enabled manual pruning of outliers and recalibration of Euler angles. This yielded a refined dataset of 12,777 high-confidence particles. For final reconstruction, particle coordinates were re-centered on a shorter segment encompassing only three dimeric units and re-extracted with a smaller box size of 64 pixels at full pixel size. The resulting particles were subjected to 3D auto-refinement, followed by separate refinements for the Healthy subset of 3,004 particles and the Alport syndrome subset of 9,773 particles. This yielded density maps at resolutions of 11□Å and 9.5□Å, respectively, as estimated by the gold-standard Fourier shell correlation (FSC = 0.143) criterion.

### Subtomogram averaging of the slit diaphragm

For the ectodomain of the slit diaphragm, 16 high-quality tomograms in the first dataset were selected for manual particle picking in IMOD 4.11.3. Given the relatively low density of the slit diaphragm, the center of particles was chosen on the podocyte membrane, ensuring suitability for subsequent averaging. The selected coordinates were sampled using the *relion_helix_toolbox* in RELION/3.1 ^59^, with a sampling interval of 120 Å as measured in IMOD, yielding 670 particles. Particle extraction was performed in Warp 1.0.9 at binning factor 2 with a box size of 196 pixels, followed by averaging in RELION. During the first classification, global classification was applied with restrictions on the psi and tilt angles, and the reference was a rectangle resembling the cell membrane, generated by MATLAB. The classification yielded 612 particles with strong slit diaphragm densities on the extracellular side of the cell membrane. This particle selection process was repeated for an additional 107 tomograms containing the slit diaphragm, comprising 33 tomograms from dataset 2, 29 tomograms from dataset 3, and 45 tomograms from dataset 4. Because the Euler angles of the slit diaphragm are relatively fixed between podocytes, the newly selected particles were manually adjusted to approximate correct orientations using the ArtiaX plugin in ChimeraX-1.10.1.

During the classification of the second dataset, the refined 3D map generated from the first dataset was employed as a reference. High-quality class averages from the second dataset were subsequently merged with the first dataset, followed by iterative Euler angle refinement. Following global orientation optimization of all particles, local refinement was performed, which significantly enhanced the density features of the ectodomain. The particles were then repositioned using TOM toolbox ^60^ in MATLAB to align their centers with the midpoint of the slit diaphragm between the two cell membranes, followed by re-extraction at binning factor 2 with a box size of 196 pixels.

For the third and fourth datasets, particles were classified and their coordinates were merged with those from the first two datasets for refinement. This process significantly improved the density of the slit diaphragm. The refined particle coordinates were then centered on the slit diaphragm, and particles were re-extracted at full pixel size with a box size of 280 pixels, followed by local refinement with restricted Euler angles. To further remove suboptimal particles, 3D classification was performed, resulting in a final set of 4,052 particles. These particles were subsequently subjected to Euler angle refinement in ArtiaX and alignment in RELION/3.1.

For symmetry expansion, particle coordinates were flipped along the X-axis, and refinement was performed with tilt and psi angle restrictions. Then, all particles were merged, yielding 8,104 particles, which were further classified to remove poor-quality particles, ultimately retaining 5,626 particles with a resolution of 22.1 Å. Finally, the Healthy and Alport syndrome datasets were processed separately, and a slit mask encompassing a single repeat unit, designated as slit mask 3, was applied during final refinement, achieving resolutions of 23.6 Å and 26.2 Å, respectively.

For the intracellular domain of the slit diaphragm, the aligned ectodomain of the slit diaphragm was centered on both cell membranes, with the Euler angles of the particles from one side flipped to align with the orientation of the particles from another side. After merging the particles from both sides, particle extraction was performed in Warp 1.0.9 at full pixel size with a box size of 280 pixels. The tilt and psi angles of the particles were constrained, and a rectangular mask was applied to the ectodomain. Local refinement was then performed in RELION/3.1. Subsequently, three Euler angles were restricted, and a rectangular mask was applied below the membrane for local classification. A class showing a well-ordered cell membrane was selected from the results, and local refinement was performed using the same mask, yielding the final structure.

### The slit diaphragm modeling

The slit diaphragm model was built using rigid body fitting in ChimeraX-1.10.1. Initially, the homologous structure SYG1-SYG2 from *C*. elegans (PDB: 4OFY) was used to determine the correct positioning of the Nephrin-Neph1 interaction site within the map. The individual Ig-like domains were then fitted into the map according to the orientation of the internal structure. Once the Nephrin-Neph1 heterodimer was properly positioned, real space refinement was performed in Coot-0.9.6.1 to prevent clashes. The final heterodimer model was further manipulated by rotation and translation to generate the tetramer structure, which was rendered in ChimeraX.

### General steady-state albumin advection-diffusion formulation

Albumin transport through GBM throats, the slit diaphragm, and the two bulk-fluid regions was described using a common steady one-dimensional advection-diffusion formulation. For a one-dimensional transport element, the albumin flux density *J*_alb_ per unit albumin-accessible area was written as

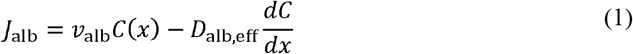

where *C*(*x*). is the equivalent albumin concentration along the transport direction, *v*_alb_ is the effective axial convective velocity of albumin, and *D*_alb,eff_ is the effective albumin diffusion coefficient.

For a transport element of length *L*, the Péclet number Pe was defined as

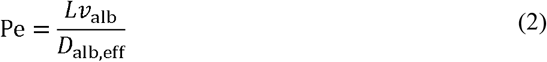

representing the ratio of albumin advection over the transport length to diffusion over the same distance. A larger Pe therefore indicates a more convection-dominated throat-transport regime. With *x* defined along the local solvent-flow direction and boundary conditions *C*(0)=*C*_in_ and *C*_(*L*)_=*C*_out_, the steady-state solution of Eq. (1) gives

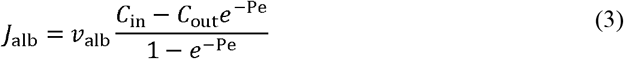

The corresponding albumin flow rate Q_alb_ was obtained by multiplying the flux density by the albumin-accessible area:

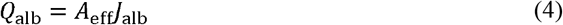

The GBM, slit diaphragm, and bulk-layer calculations below used this common solution with layer-specific definitions of *A*_eff_, *v*_alb_, *D*_eff_, *L*, and the corresponding boundary conditions.

### Albumin sieving model for the slit diaphragm

The slit diaphragm was modeled as a single transport interface rather than a pore-throat network because the slit diaphragm architecture was resolved at a more explicit structural level. To match this explicit slit diaphragm geometry, albumin was represented using a fitted ellipsoidal molecular geometry rather than the spherical approximation used in the GBM pore-throat model. This explicit geometric representation was used to determine the albumin-accessible slit diaphragm area, which was then coupled to the common one-dimensional transport calculation.

For the slit diaphragm layer, Eq. (1) was applied with *J*_alb_=*J*_SD_, *v*_alb_, =*v*_alb,SD_, and *D*_alb,eff_=*D*_*alb*_ · *D*_*alb*_ is the free diffusion coefficient of albumin, calculated using the Stokes–Einstein relation and listed in Extended Data Table 1.

The mean solvent velocity through the slit diaphragm interface was defined from the conserved water flow rate and the water-accessible slit diaphragm area:

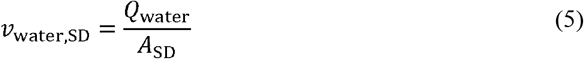

To approximate the higher local solvent velocity sampled by albumin centers within the accessible region, the effective albumin axial convective velocity was written as

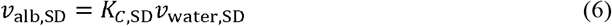

The SD Péclet number was defined as the ratio of advective transport across the slit diaphragm layer to diffusive transport:

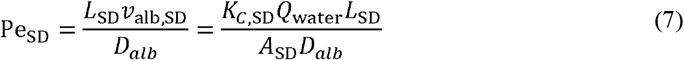

where *L*_SD_ is the effective slit diaphragm protein thickness, as listed in Extended Data Table 1.

Applying Eqs. (3) and (4) across the slit diaphragm layer, with *C*_in_=*C*_SD,in_, *C*_out_=*C*_SD,out_,*A*_eff_=*A*_eff,SD_ and Eqs. (6) and (7) gives

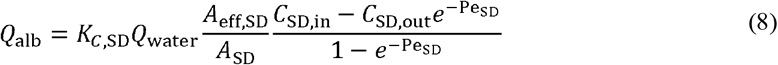

The downstream side of the slit diaphragm was treated as a well-mixed reservoir, so the outlet-equivalent concentration satisfies the mixing balance:

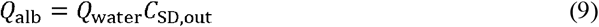

Combining Eqs. (8) and (9) for *Q*_alb_, and dividing by *Q*_water_, gives

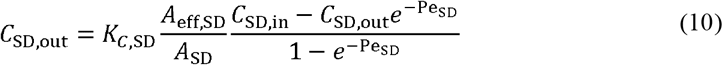

The slit diaphragm sieving coefficient was defined as

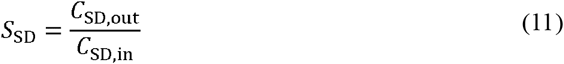

Dividing Eq. (10) by *C*_SD,in_ gives

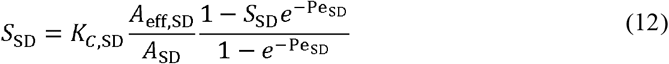

The albumin-accessible area fraction was then defined as

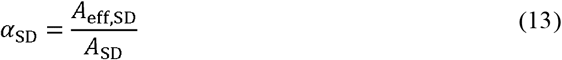

And under the conditions analyzed in this study, Pe_SD_ « 1. With 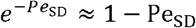, Eq. (12) becomes

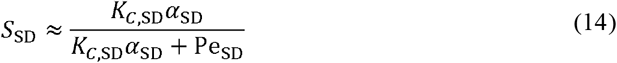

In this expression, Pe_SD_ was defined using the albumin convective velocity *v*_alb,SD_ = *K*_*c*,SD_ *v*_water,SD_ Equivalently, if the slit diaphragm Péclet number is written using the mean solvent velocity,

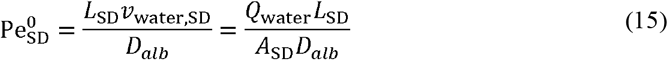

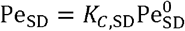, and the same approximation becomes

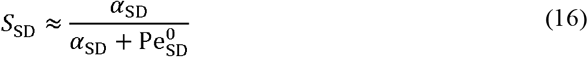

Thus, the constant convective correction factor *K*_*C*,SD_ does not change the simplified low-Pe sieving form when the Péclet number is expressed using the mean solvent velocity. The simplified slit diaphragm sieving expression therefore highlights the competition between geometric accessibility, represented by α_SD_, and convective transport relative to diffusion, represented by 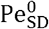.

### Orientation-Bounded Estimation of slit diaphragm Accessible Area and Sieving Coefficient

The structural input *A*_eff,SD_ in the slit diaphragm transport model was estimated from the slit diaphragm pore geometry and an ellipsoidal albumin geometry. To define the reference slit diaphragm pore geometry, the slit diaphragm protein structure was projected onto two-dimensional planes from different viewing directions, and the major slit diaphragm pore was approximated as a hexagonal opening.

Because the slit diaphragm proteins are anchored to opposing podocyte membranes, changes in slit diaphragm width were assumed to stretch or compress the pore geometry. Therefore, for each measured slit diaphragm width, the polygonal pore shape was deformed by adjusting the pore height and internal angles while keeping the lengths of the surrounding protein segments approximately constant. The measured slit diaphragm width distribution was fitted as a continuous probability distribution *P*(*W*), which was used to weight the width-dependent pore geometries.

Because direct information on the three-dimensional orientation distribution of albumin during slit diaphragm passage is not available, slit diaphragm accessibility was evaluated using two limiting orientation scenarios. In the lower-bound scenario, albumin was assumed to encounter the slit diaphragm pore with a uniformly random three-dimensional orientation, so accessibility was determined by the instantaneous orientation at the pore entrance. In the upper-bound scenario, albumin was assumed to remain near the slit diaphragm pore long enough to sample available orientations and adopt the most favorable three-dimensional orientation for passage at each slit diaphragm width.

For each slit diaphragm width *w* and albumin three-dimensional orientation Ω, *A*_eff,SD_(*w*,Ω) was defined as the area available to the center of the albumin footprint while the entire albumin footprint remained inside the slit diaphragm pore. This center-accessible region was computed as a geometric erosion of the pore by the albumin footprint:

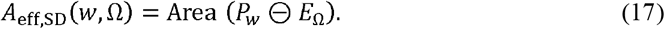

where *P*_*w*_ is the deformed slit diaphragm pore geometry at width _*w*_, *E*_Ω_ is the albumin footprint on the slit diaphragm pore plane for orientation Ω, and ⊖ denotes geometric erosion. In physical terms, *P*_*w*_ ⊖ *E*_Ω_ is the set of all albumin-center positions for which the albumin footprint can be placed without crossing the protein boundary. If the albumin footprint could not be fully accommodated within the pore, *A*_eff,SD_(*w*,Ω) was set to 0.

The same width-weighted integration was then applied to both orientation scenarios. For each slit diaphragm width, Ā_eff,SD_(*w*) denotes the orientation-processed accessible area: the mean of A_eff,SD_(*w*, Ω) over uniformly random three-dimensional orientations for the lower-bound calculation, or the maximum value of A_eff,SD_(*w*, Ω) over sampled three-dimensional orientations for the upper-bound calculation. The integrated accessible area for each scenario was calculated as:

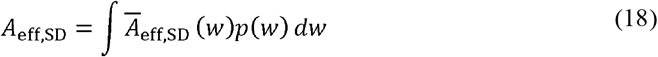

The corresponding lower- and upper-bound accessible area fractions, 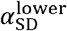 and 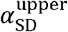, were obtained using the area-fraction definition in Eq. (13). These two values were substituted separately into the slit diaphragm sieving relation in Eq. (16) to obtain 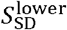 and 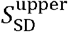. The final slit diaphragm sieving coefficient used in the serial GFB transport model was then estimated as the midpoint of these two limiting sieving coefficients:

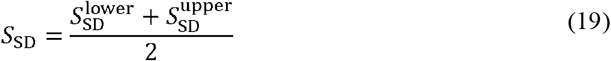

This formulation treats random orientation and best orientation as lower and upper accessibility bounds, respectively, and avoids assigning a single fixed albumin orientation during slit diaphragm passage.

### Pore network modeling of the lamina densa

3D segmentation of the GBM was performed on denoised cryo-ET data using Avizo 2024 software (Thermo Fisher Scientific). Initial pore segmentation was achieved via the Interactive Thresholding module based on voxel grayscale intensity. To specifically isolate the lamina densa, a 3D mask was applied to constrain the region of interest, enabling the independent segmentation of the fibrillar and pore phases. To further elucidate molecular transport properties, a pore network model (PNM) was constructed. The Generate Pore Network Model module was subsequently employed to extract key parameters of the resulting pore-throat network representation of the lamina densa, including the number, radii, and topological connectivity of pores and throats.

### Identification of inlet- and outlet-side surface pores

To impose inlet- and outlet-side boundary conditions in the pore-throat network model, surface pores were identified along the assigned filtration direction. Because reconstructed GBM boundaries were irregular rather than ideal planes, pores were not selected simply by their minimum or maximum axial coordinates; such a criterion could incorrectly include side-wall pores, pores in local depressions, or pores affected by cropping. We therefore used a surface-detection procedure combining alpha-shape boundary reconstruction, ray-based visibility testing, and geometric filtering ^61^.

Briefly, an alpha-shape envelope was constructed from pore-center coordinates to approximate the outer boundary of the reconstructed pore space. The alpha parameter was estimated from local pore spacing, with convex-hull reconstruction used as a fallback when alpha-shape construction failed. Candidate pores were then evaluated in a filtration-axis coordinate system, where the two axial end surfaces were represented by the *u*_min_, and *u*_max_ sides. For each candidate pore, rays were cast from points outside the corresponding axial end surface toward the pore center, and pores with unobstructed ray paths were classified as directly visible from that surface.

These visible candidates were further refined by geometric filters that removed pores associated with lateral boundaries, local surface depressions, isolated surface artifacts, or cropping edges. Retained candidates were also required to lie within a radius-scaled distance from the corresponding GBM end surface. After these filtering steps, retained *u*_min_-side pores were defined as inlet-side surface pores, and retained *u*_max_-side pores were defined as outlet-side surface pores. The resulting classification was written to the pore-classification table and used for boundary-condition assignment in subsequent transport calculations.

### Pore-throat transport model for GBM albumin sieving

The three-dimensional void space of the GBM lamina densa was represented as a pore-throat network, in which larger connected void compartments were treated as pore nodes and constricted interconnecting segments were treated as throat edges. Inlet and outlet nodes were defined according to the filtration direction, corresponding to the capillary-facing and Bowman’s-space-facing boundaries of each reconstructed GBM volume. The model assumed steady state, such that the pressure field and albumin concentration field within the GBM did not vary with time.

At steady state, each internal pore node satisfies conservation of solvent and albumin flow rates:

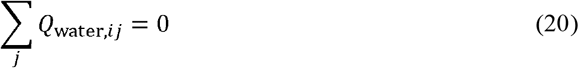

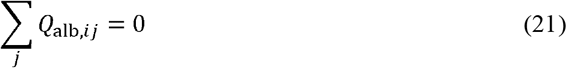

where *Q*_water,*ij*_ and *Q*_alb,*ij*_ denote the solvent volumetric flow rate and albumin flow rate, respectively, through the throat connecting pore *i*to neighboring pore *j*. The summation is over all throats connected to pore *i*. The albumin conservation equation assumes no albumin adsorption or degradation within internal pores and throats under the steady-state formulation.

Each throat was approximated as a cylindrical hydraulic resistor. The volumetric solvent flow rate through a throat was calculated using Poiseuille’s law:

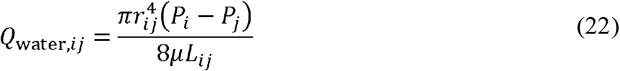

where *r*_*ij*_ is the effective throat radius, *L*_*ij*_ is the throat length, *P*_*i*_ − *P*_*j*_ is the pressure difference between the connected pores, and μ is the dynamic viscosity at 37 ° C, which is listed in Extended Data Table 1.

A preliminary hydraulic-resistance estimate comparing GBM resistance derived from simulated Healthy pore-throat network flow rates with slit diaphragm resistance approximated by a rectangular slit channel model indicated that the hydraulic pressure drop was overwhelmingly dominated by GBM permeation. Therefore, the pressures of inlet and outlet pore nodes used in the hydraulic calculation were fixed at the corresponding plasma and Bowman’s-space pressures listed in Extended Data Table 1. Pressures were converted to SI units before solving the linear systems.

Substituting Eq. (22) into the solvent-conservation equations in Eq. (20) yielded a linear system in the unknown internal pore pressures. Once the pressure field was obtained, *Q*_water,*ij*_ and the corresponding mean axial solvent velocity in each throat were determined.

Because GBM throat radii are comparable to the hydrated size of albumin, albumin transport was calculated only through the geometrically accessible portion of each throat. Albumin was approximated as a sphere with a baseline hydrated radius *r*_alb_, as listed in Extended Data Table 1. Throats with *r*_*ij*_ ≤*r*_alb_ were classified as albumin-inaccessible, whereas throats with *r*_*ij*_ > *r*_alb_ were defined as albumin-accessible. Only albumin-accessible throats and pores belonging to components connected to both the inlet and outlet boundaries were retained for trans-GBM albumin transport calculations.

For each retained throat, the solute-to-throat size ratio was defined

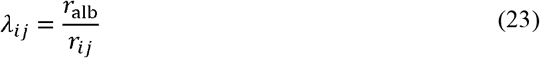

The cross-sectional area accessible to the center of an albumin molecule was

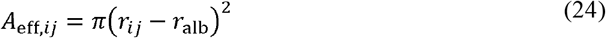

This effective-area formulation accounts for the exclusion of albumin centers from the region within one albumin radius of the throat wall. Equivalently, albumin was represented as a point-like solute moving within the accessible central region of each throat. This formulation allowed a single equivalent albumin concentration variable to be assigned to each pore node without introducing separate pore-throat partition-equilibrium transformations at individual junctions.

The water-accessible cross-sectional area of throat *ij* was 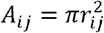. The mean axial solvent velocity was therefore

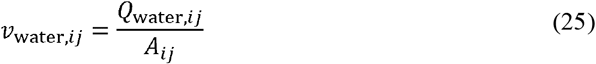

Although *v*_water,*ij*_ characterizes the cross-sectionally averaged solvent motion, a finite-sized albumin molecule does not necessarily move at this velocity or retain its free-solution diffusivity within a confined throat. Geometric accessibility alone does not account for the hydrodynamic interaction between a finite-sized albumin molecule and the cylindrical throat wall. This interaction modifies both the Brownian diffusional mobility of albumin and its effective convective velocity relative to the cross-sectionally averaged solvent velocity. We therefore introduced two dimensionless transport factors, *K*_*D,ij*_ and *K*_*C,ij*_, both determined by the solute-to-throat size ratio λ_*ij*_. The diffusive hindrance factor *K*_*D,ij*_, representing the ratio of the restricted albumin diffusion coefficient within the accessible region to the free-solution diffusion coefficient, and the convective factor *K*_*C,ij*_, representing the ratio of the effective albumin convective velocity to the cross-sectionally averaged solvent velocity, were calculated as functions of λ_*ij*_. using the cylindrical-pore approximations summarized by Dechadilok and Deen ^19^. For 0≤_*ij*_ ≤ 0.95 the diffusive hindrance factor was:

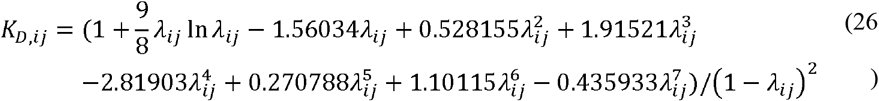

When λ_*ij*_ =0, the term λ_*ij*_ ln(λ_*ij*_) was evaluated using its limiting value of zero.

For the near-occlusion range 0.95 < λ_*ij*_ < 1, the corresponding asymptotic approximation was used:

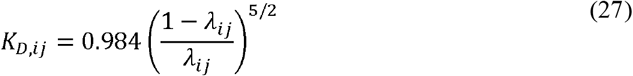

The effective albumin diffusion coefficient within throat *ij* was then

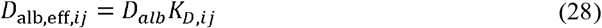

The convective hindrance factor was calculated from the same size ratio as

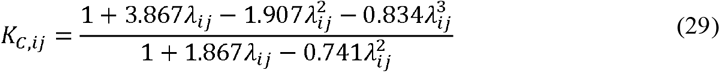

The effective axial convective velocity of albumin was therefore

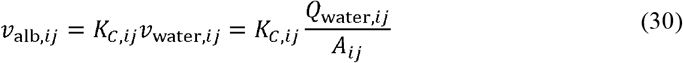

For throat *ij*, the layer-specific Péclet number was

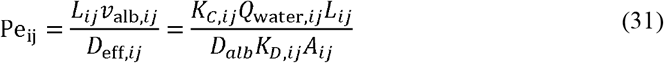

To formulate the network-level albumin transport problem, all equivalent albumin concentrations were normalized by the concentration entering the GBM. The inlet boundary condition was therefore set to *C*_GBM.in_ = 1, representing a relative GBM-inlet concentration rather than an absolute concentration of 1. All albumin-accessible inlet pores were assigned this normalized concentration. At the downstream boundary, albumin leaving different outlet pores was assumed to enter a common well-mixed reservoir with an unknown equivalent concentration *C*_GBM.out_.

With these boundary conditions, the throat-specific definitions in Eqs. (23)–(31) were substituted into the common analytical solution in Eqs. (3) and (4), using *C*_in_ = *C*_*i*_, *C*_out_ = *C*_*j*_, *L* = *Lij*, and *A*_eff_=*A*_eff,*ij*_. The resulting albumin flow rate through each retained throat was a linear function of the equivalent concentrations at its two connected pores. Applying the albumin-conservation equation in Eq. (21) at every internal pore and imposing the global albumin-flow balance at the common outlet produced a coupled linear system for the internal pore concentrations and *C*_GBM,out_. Solving this system yielded the complete GBM concentration field and the corresponding outlet-equivalent albumin concentration.

For an individual reconstructed sample, the GBM sieving coefficient was defined in the conventional concentration-ratio form:

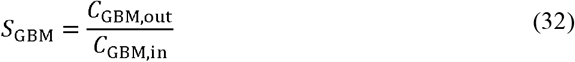

Because *C*_GBM,in_ was normalized to 1, *S*_GMB_was numerically equal to the solved outlet-equivalent albumin concentration *C*_GBM,out_. If the albumin-accessible throat network did not contain a continuous connected pathway from inlet to outlet, the albumin flow rate was assigned as 0, representing a physically albumin-blocked network result.

### Subsample segmentation

After the filtration direction was determined for each original GBM reconstruction, local subsamples were generated at two spatial scales. In all subsamples, pores were retained only when their centers were located within the selected region, and throats were retained only when both connected pores were located within that region.

For real-sample albumin sieving-coefficient calculations, a single relatively large representative subsample was selected from each original reconstruction. The selected region was chosen to preserve a locally complete pore-throat network while avoiding abrupt thickness variation along the filtration direction (Extended Data Fig. 16A). These large subsamples were used for real-sample structural analysis and albumin sieving-coefficient calculation.

For thickness-dependent structural statistics and fitted-network generation, each original reconstruction was further divided into smaller local subsamples with relatively uniform thickness and balanced in-plane dimensions. The shorter in-plane dimension of each sample was used as the reference length, and approximately square regions were cropped along the longer in-plane direction, with preference given to regions showing smaller thickness variation (Extended Data Fig. 16B). When neighboring small subsamples overlapped spatially, pores and throats in the overlapping region were retained in each corresponding subsample. Thus, small subsamples were treated as local structural sampling units rather than mutually exclusive volume partitions.

### Thickness measurement, tilt correction, and thickness sampling

GBM thickness was calculated for each subsample according to its assigned filtration direction. Because the sectioning plane was not always perpendicular to the physiological filtration direction, thickness measured directly along the imaging axis could overestimate or underestimate the true trans-GBM distance. Therefore, the two GBM boundary surfaces were used to estimate the local surface orientation and filtration axis, and the tilt angle θ relative to the imaging plane was calculated. When the surface-orientation estimate was considered reliable and θ exceeded 25°, the thickness measured along the filtration axis was corrected as *T*_corrected_ = *T*_axiz_ cosθ. When θ did not exceed 25°, the along-axis thickness was used directly. For samples in which the tilt estimate was unstable and produced an evidently unreasonable corrected thickness, the thickness value was manually reviewed using local thickness quality-control measurements, and a consensus thickness was assigned. The quality-control-reviewed thickness values were used for real-sample stratification, thickness binning, and fitted-network generation.

For fitted-network generation, Healthy and Alport syndrome thickness distributions were estimated separately from the quality-control-reviewed thicknesses of small subsamples. To avoid imposing a strong parametric assumption on the shape of the thickness distribution, kernel density estimation (KDE) was used to describe the Healthy and Alport syndrome thickness distributions _62_.

### Thickness-dependent structural parameter fitting for pore-throat network generation

To generate fitted GBM pore-throat networks, structural parameters were quantified from small Healthy and Alport syndrome subsamples across thickness conditions. Each small subsample was assigned to a thickness bin according to its mean GBM thickness. Object-level parameters, including pore radius, throat radius, throat length, and pore degree, were analyzed using 10 nm thickness bins by default, whereas subsample-level pore and throat number densities, calculated as the number of pores or throats divided by subsample volume, were analyzed using 20 nm bins to ensure sufficient samples for distribution fitting. Sparse bins were merged with adjacent bins or smoothed using nearby thickness-bin fits when necessary. For Alport syndrome samples, high-thickness bins above 160 nm were pooled because sample numbers were limited and structural parameters showed weaker thickness dependence in this range (Extended Data Fig. 17).

Across these two parameter levels, continuous positive-valued parameters, including pore number density, throat number density, pore radius, throat radius, and throat length, were fitted separately for Healthy and Alport syndrome within each thickness bin using candidate distributions including lognormal, gamma, normal, and Gaussian mixture models. Model selection was based primarily on the Bayesian information criterion, with the Kolmogorov-Smirnov statistic used as a secondary goodness-of-fit criterion ^63–65^. When a normal distribution performed similarly to a positive-valued distribution, the positive-valued distribution was preferred to avoid sampling negative values during network generation. Pore degree was treated as a discrete parameter and sampled from empirical probability distributions constructed within each thickness bin.

To preserve structural correlations during fitted-network generation, pairwise relationships among selected object-level parameters were fitted within each thickness bin. These relationships included those between throat radius and adjacent pore radii and between throat radius and throat length. The fitted pairwise relationships were used during network generation to reduce unrealistic combinations of independently sampled parameters and to better retain the local geometric structure observed in reconstructed networks (Extended Data Fig. 11).

### Fitted GBM pore-throat network generation

To estimate albumin sieving across a broader range of GBM geometries than directly reconstructed samples, fitted GBM pore-throat networks were generated using condition-specific thickness distributions and thickness-dependent structural-parameter fits derived separately from Healthy and Alport syndrome small subsamples. Each fitted network was represented as a standardized cuboid GBM volume with fixed in-plane dimensions *L* × *L*and thickness *T*. The in-plane dimension *L* was chosen to match the lateral extent of the analyzed pore-throat networks in real reconstructed samples and is listed in Extended Data Table 1. The thickness T was sampled from the corresponding Healthy or Alport syndrome KDE-fitted thickness distribution, with values outside the predefined interpretable range redrawn by rejection sampling. A uniform-thickness mode was used, such that no spatially varying thickness field was introduced within an individual generated network.

After thickness assignment, the generation domain was divided into 50 nm in-plane grid cells. For each grid cell, local pore density was sampled from the corresponding thickness-binned pore-density distribution, and the expected pore number was calculated from the sampled density and local cell volume. Realized pore numbers were generated by stochastic rounding, and pore centers were placed uniformly within each grid cell across the full network thickness. Pore radii were then sampled from the fitted pore-radius distribution for the corresponding thickness bin, with a lower radius bound of 0.5 nm.

Throat connections were generated after pore placement and pore-radius assignment. Candidate pore pairs were identified within a local neighborhood using a KD-tree search, with the search distance listed in Extended Data Table 1 ^66^. Connections were selected by weighted sampling to match the thickness-dependent throat-density, throat-length, and pore-degree statistics measured from reconstructed samples. In this procedure, throat length was determined by the center-to-center distance of the selected pore pair; therefore, the fitted throat-length distribution and sampled pore-degree targets guided candidate selection and connection propensity rather than assigning these properties independently.

Throat radii were assigned after throat connections were determined. Radii were sampled from the corresponding thickness-binned throat-radius distribution and assigned using weighted matching that incorporated fitted relationships between throat radius, adjacent pore radius, and throat length. This procedure allowed generated networks to reproduce thickness-dependent marginal distributions while retaining selected structural correlations observed in reconstructed networks.

### Integrated sieving coefficient calculation

For each fitted GBM network ensemble, we calculated an integrated albumin sieving coefficient to estimate the effective albumin permeability of intact GBM. Each generated network was analyzed using the same pore-throat transport model applied to reconstructed samples.

The conventional sieving coefficient is a concentration ratio. For a well-mixed outlet of GBM network *k*, the outlet-equivalent albumin concentration is the albumin flow rate divided by the solvent flow rate:

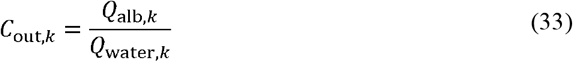

Therefore, the concentration-ratio definition can be written equivalently as a flow-rate ratio normalized by the inlet concentration:

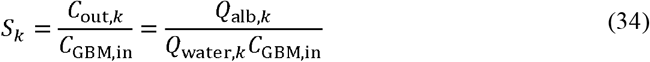

For an ensemble of fitted networks, the integrated coefficient was defined by first summing the albumin and water flow rates across networks and then taking the same outlet-mixing ratio:

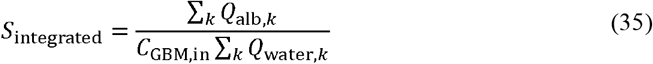

Albumin concentration was normalized by the concentration entering the GBM. Therefore, *C*_GBM,in_ = 1 for all reconstructed and fitted networks, and Eq. (35) was numerically equivalent to 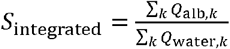. This definition is therefore a water-flow-weighted effective sieving coefficient rather than an arithmetic mean of individual network sieving coefficients. Networks with no connected albumin-accessible pathway from inlet to outlet were retained as physically disconnected cases, with *Q*_alb,*k*_ *=* 0.

### Parameter replacement and sieving-coefficient sensitivity analysis

To assess how selected structural parameter groups influenced model-predicted albumin sieving, parameter-distribution replacement analyses were performed using the fitted-network generation workflow. For each replacement direction, baseline and replacement networks were generated using the same sampled thickness table and paired by run index. This pairing controlled for differences in sampled GBM thickness between baseline and replacement conditions.

Two replacement directions were tested. In the Alport syndrome-to-Healthy direction, fitted Alport syndrome networks were used as the baseline, and selected Alport syndrome parameter distributions were replaced by the corresponding Healthy distributions. In the Healthy-to-Alport syndrome direction, fitted Healthy networks were used as the baseline, and selected Healthy parameter distributions were replaced by the corresponding Alport syndrome distributions. Because Healthy and Alport syndrome thickness distributions differed substantially, replacement-source distributions were pooled across thickness bins using sample-number weighting before replacement. This allowed structural parameter effects to be assessed without also imposing the source condition’s thickness distribution.

The replacement conditions focused on throat-related structural features and pore connectivity because, in the GBM pore-throat model, hydraulic resistance and albumin accessibility are controlled primarily by the constricted throats and by how throats connect the pore space. Therefore, throat radius, throat number density, throat length, and pore degree were selected as the main replacement targets. Because fitted-network generation preserved geometric correlations between throats and neighboring pores, replacing some throat-related parameters, such as throat radius or throat density, could also alter pore-related summary statistics indirectly. Each replacement network was then passed through the same complete transport-analysis workflow as its paired baseline network. The resulting sieving-coefficient distributions and integrated sieving coefficients were compared with the paired baseline condition (Extended Data Fig. 13A–B).

To verify that each replacement altered the intended network features, generated structural-parameter distributions were summarized under each condition. These summary metrics included pore density, mean pore radius, throat density, mean throat radius, mean throat length, and mean pore degree. Pore-related quantities were included in this verification step because throat generation is geometrically coupled to pore abundance and pore size, even when the primary replacement targeted throat-related parameters (Extended Data Fig. 13C–D).

### Effective-radius approximation for electrostatic exclusion in GBM transport

To estimate the potential effect of electrostatic exclusion on albumin transport through the GBM, electrostatic repulsion between negatively charged albumin and negatively charged GBM structure was represented as an additional exclusion distance. In water, this exclusion distance was approximated by the Bjerrum length, *l*_*B*_, defined as

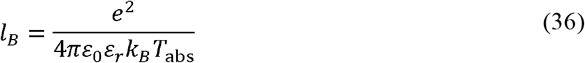

Where *e* is the elementary charge, *ε*_0_ is the vacuum permittivity, *ε*_*r*_ is the relative permittivity of water at 37°C, *k*_*B*_ is the Boltzmann constant, and *T*_abs_ is the absolute temperature. The value of *l*_*B*_ used in this study is listed in Extended Data Table 1.

The effective albumin radius used for the electrostatic-exclusion calculation was then defined as

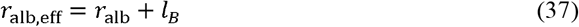

This increased effective radius reduces the GBM pore or throat space accessible to albumin centers. *r*_alb,eff_ was used only to determine throat accessibility and the albumin-center-accessible area. The hydrated albumin radius of 3.55 nm was retained when calculating *K*_*D*_, *K*_*C*_, and the associated hydrodynamic transport coefficients.

### Bulk-layer transport in the serial four-layer model

The two bulk regions in the serial filtration model, corresponding to the plasma-to-GBM boundary layer and the GBM-to-slit diaphragm cleft, were modeled as one-dimensional steady advection-diffusion layers without geometric sieving.

Under steady-state serial transport, the solvent flow rate was conserved across the bulk layers, GBM, and slit diaphragm layer:

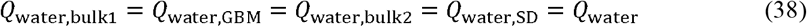

The albumin flow rate was likewise conserved:

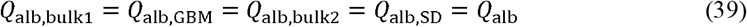

In the bulk layers, the albumin-accessible area was equal to the water-accessible area:

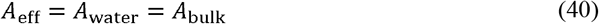

Thus, the bulk layers did not impose additional size exclusion but instead described concentration polarization under conserved water and albumin flow rates.

For the bulk layers, the common transport equation in Eq. (1) was applied with *v*_alb_ = *v*_bulk_ and *D*_alb,eff_ = *D*_*alb*_. Because *A*_eff_ = *A*_bulk_, the albumin flux density was

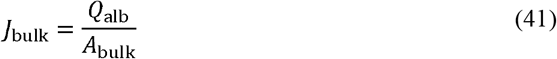

and the mean convective velocity was

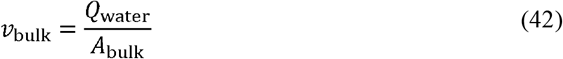

Their ratio is therefore independent of the bulk-layer area:

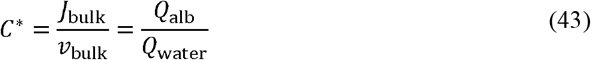

The quantity *C*^*^ is the conserved albumin-to-water flow-rate ratio shared by all layers in the serial model. It can be interpreted as the equivalent concentration carried by the filtrate after complete local mixing. The same GBM-determined *Q*_water_ was used in both bulk layers as the conserved solvent volumetric flow rate.

Substituting Eq. (43) into the common transport equation gives

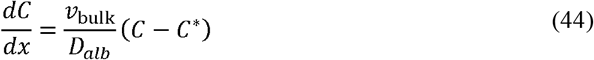

With *x* defined along the filtration direction, the analytical solution for the outlet concentration is

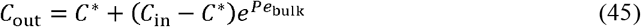

Where

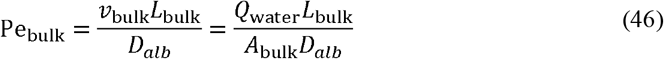

Here, *L*_bulk_ is the effective bulk-layer thickness listed in Extended Data Table 1.

The transfer relation in Eq. (45) was applied to both bulk layers, using the corresponding inlet and outlet boundary concentrations for the plasma-to-GBM boundary layer and the GBM-to-slit diaphragm cleft. Together with the effective GBM and slit diaphragm sieving relations, these equations formed a self-consistent serial-layer system in which *C*^*\**^ was solved under conserved water and albumin flow rates. This formulation keeps the original concentration-ratio definition of sieving while using flow-rate conservation to couple the layers.

For the parameter values used in this study, Pe_bulk_ « 1, and concentration polarization across the bulk layers remained at a very low level.

### Stochastic fiber network model

To evaluate whether a continuous stochastic fiber-network model could account for albumin sieving by the GBM, we approximated the GBM as a homogeneous matrix of randomly distributed cylindrical fibers. The fiber radius was denoted by *r*_*f*_, the fiber length density per unit volume by *L*_*f*_, and the fiber volume fraction by *V*_*f*_. The parameter values used in this analysis are listed in Extended Data Table 1. For long cylindrical fibers,

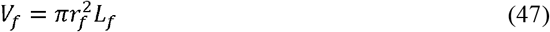

The stochastic fiber-network model does not represent the GBM as a set of regular cylindrical pores. Instead, sieving is estimated from the probability that albumin is not geometrically excluded by randomly arranged fibers. Specifically, when a spherical solute with radius *r*_*s*_ is randomly placed in the fiber matrix, the probability that its center does not geometrically overlap with any fiber is defined as the partition coefficient, *Φ*. This coefficient represents the fraction of water-accessible space that is accessible to the center of albumin.

Following the statistical insertion framework of the stochastic fiber-network model, we treated this geometric exclusion process as a two-step insertion process:

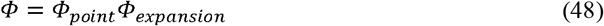

where *Φ*_*point*_ represents the probability that a point solute can enter the random fiber matrix, and *Φ*_*expansion*_ represents the probability that this point solute remains accessible after being expanded to a finite-radius solute^67^.

First, we treated the solute as a point particle and approximated random fiber-exclusion events as a Poisson process. In this case, the fiber-occupied volume fraction is *V*_*f*_, and the zero-collision probability that the point solute does not fall within the fiber-occupied region is:

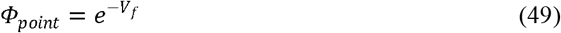

The point solute was then expanded from *r*_*s*_ = 0 to a finite radius *r*_*s*_.

For a solute with radius *r*_*s*_, the exclusion density for the solute center increases relative to the point-solute case. The exclusion density for a finite-radius solute can be written as:

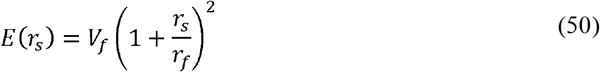

whereas for a point solute:

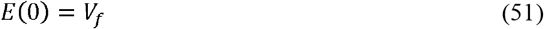

Thus, the additional excluded volume fraction introduced by the finite solute radius, relative to a point solute, is:

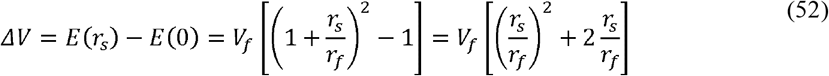

Because this additional excluded volume must be accommodated within the remaining void space, 1−*V*_*f*_, the corresponding expansion probability is:

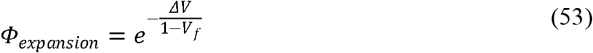

Thus, the albumin partition coefficient used in the stochastic fiber-network model is:

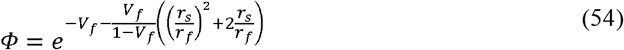

In classical stochastic fiber-based estimates of GBM sieving, a high-Peclet-number, convection-dominated approximation is commonly used. Under this approximation, albumin sieving is governed primarily by whether albumin can be convectively carried through the fiber matrix with water, whereas local diffusive contributions are simplified. We defined the reflection coefficient, *σ*, as the fraction of albumin rejected relative to solvent flow; *σ*= 0 indicates that the solute moves freely with water, whereas *σ*= 1 indicates complete convective rejection. Based on the hydraulic-radius approximation, convective solute transmission through a fibrous network requires both geometric accessibility and effective solvent drag, so the solvent-drag reflection coefficient is approximated as scaling with the square of the inaccessible fraction, 1− *Φ*^68^:

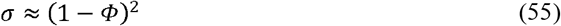

Under this approximation, the fraction of albumin transmitted with water can be written as:

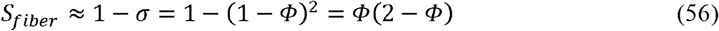

We used this expression in two directions.

First, for a specified target GBM sieving coefficient, *S*_*target*_, the equivalent fiber volume fraction required by the stochastic fiber-network model was obtained by numerically solving:

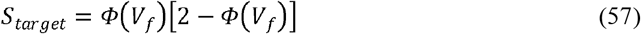

Second, for specified values of *v*_*f*_, *r*_*f*_, and *r*_*s*_, we directly calculated *Φ* and *S*_*fiber*_ to estimate the albumin sieving coefficient predicted by the stochastic fiber-network model.

## Notes

### Competing Interest Statement

The authors have declared no competing interest.

### Summary of Updates

We have updated several statements to make the text accurate.

## References

1 Haraldsson, B., Nyström, J. & Deen, W. M. Properties of the glomerular barrier and mechanisms of proteinuria. Physiol Rev 88, 451–487 (2008). 10.1152/physrev.00055.2006

2 Grahammer, F., Schell, C. & Huber, T. B. The podocyte slit diaphragm--from a thin grey line to a complex signalling hub. Nat Rev Nephrol 9, 587–598 (2013). 10.1038/nrneph.2013.169

3 Greka, A. & Mundel, P. Cell biology and pathology of podocytes. Annu Rev Physiol 74, 299–323 (2011). 10.1146/annurev-physiol-020911-153238

4 St John, P. L. & Abrahamson, D. R. Glomerular endothelial cells and podocytes jointly synthesize laminin-1 and -11 chains. Kidney Int 60, 1037–1046 (2001).

5 Naylor, R. W., Morais, M. R. P. T. & Lennon, R. Complexities of the glomerular basement membrane. Nat Rev Nephrol 17, 112–127 (2020). 10.1038/s41581-020-0329-y

6 Suleiman, H. et al. Nanoscale protein architecture of the kidney glomerular basement membrane. eLife 2, e01149 (2013). 10.7554/eLife.01149

7 Hudson, B. G., Tryggvason, K., Sundaramoorthy, M. & Neilson, E. G. Alport’s syndrome, Goodpasture’s syndrome, and type IV collagen. N Engl J Med 348, 2543–2556 (2003).

8 Kestilä, M. et al. Positionally cloned gene for a novel glomerular protein--nephrin--is mutated in congenital nephrotic syndrome. Molecular Cell 1, 575–582 (1998).

9 Donoviel, D. B. et al. Proteinuria and perinatal lethality in mice lacking NEPH1, a novel protein with homology to NEPHRIN. Molecular and Cellular Biology 21, 4829–4836 (2001).

10 Rodewald, R. & Karnovsky, M. J. Porous substructure of the glomerular slit diaphragm in the rat and mouse. The Journal of Cell Biology 60, 423–433 (1974).

11 Wartiovaara, J. et al. Nephrin strands contribute to a porous slit diaphragm scaffold as revealed by electron tomography. J Clin Invest 114, 1475–1483 (2004).

12 Grahammer, F. et al. A flexible, multilayered protein scaffold maintains the slit in between glomerular podocytes. JCI Insight 1 (2016). 10.1172/jci.insight.86177

13 Birtasu, A. N. et al. The kidney slit diaphragm resembles a fishnet. Kidney Int (2025). 10.1016/j.kint.2025.08.028

14 Moser, D. et al. The slit diaphragm in Drosophila exhibits a bilayered, fishnet architecture. Nature Communications 16, 8741 (2025). 10.1038/s41467-025-64347-5

15 Moser, D. et al. The native human glomerulus features a slit diaphragm resembling a densely interwoven fishnet. JCI Insight 11 (2026). 10.1172/jci.insight.200658

16 Drumond, M. C. & Deen, W. M. Structural determinants of glomerular hydraulic permeability. Am J Physiol 266, F1–12 (1994).

17 Maddox, D. A., Deen, W. M. & Brenner, B. M. Glomerular Filtration. Comprehensive Physiology 1991, 545–638 (1991). 10.1002/j.2040-4603.1991.tb01569.x

18 Deen, W. M. Hindered transport of large molecules in liquid-filled pores. AIChE Journal 33, 1409–1425 (1987). 10.1002/aic.690330902

19 Dechadilok, P. & Deen, W. M. Hindrance Factors for Diffusion and Convection in Pores. Industrial & Engineering Chemistry Research 45, 6953–6959 (2006). 10.1021/ie051387n

20 Wang, H. et al. A simple and highly purified method for isolation of glomeruli from the mouse kidney. Am J Physiol Renal Physiol 317, F1217–F1223 (2019). 10.1152/ajprenal.00293.2019

21 Du, W., Yang, J. & Guo, Q. Optimized tissue sample preparation for cryo-electron tomography using serial lift-out. Biophysics Reports 11 (2025). 10.52601/bpr.2025.250035

22 Ichimura, K., Stan, R. V., Kurihara, H. & Sakai, T. Glomerular endothelial cells form diaphragms during development and pathologic conditions. J Am Soc Nephrol 19, 1463–1471 (2008). 10.1681/ASN.2007101138

23 Melo, A. A. et al. Cryo-electron tomography reveals structural insights into the membrane remodeling mode of dynamin-like EHD filaments. Nature Communications 13, 7641 (2022). 10.1038/s41467-022-35164-x

24 Vázquez-Sarandeses, E., Mikirtumov, V., Noel, J. K., Kudryashev, M. & Daumke, O. Structures of EHD2 filaments on curved membranes provide a model for caveolar neck stabilization. Nature Communications 17 (2026). 10.1038/s41467-026-76288-8

25 Patrakka, J. et al. Expression and subcellular distribution of novel glomerulus-associated proteins dendrin, ehd3, sh2d4a, plekhh2, and 2310066E14Rik. J Am Soc Nephrol 18, 689–697 (2007).

26 Chung, J.-J. et al. Single-Cell Transcriptome Profiling of the Kidney Glomerulus Identifies Key Cell Types and Reactions to Injury. J Am Soc Nephrol 31, 2341–2354 (2020). 10.1681/ASN.2020020220

27 Finch, N. C. et al. Reduced Glomerular Filtration in Diabetes Is Attributable to Loss of Density and Increased Resistance of Glomerular Endothelial Cell Fenestrations. J Am Soc Nephrol 33, 1120–1136 (2022). 10.1681/ASN.2021030294

28 Özkan, E. et al. Extracellular architecture of the SYG-1/SYG-2 adhesion complex instructs synaptogenesis. Cell 156, 482–494 (2014). 10.1016/j.cell.2014.01.004

29 Stenson, P. D. et al. The Human Gene Mutation Database (HGMD®): optimizing its use in a clinical diagnostic or research setting. Hum Genet 139, 1197–1207 (2020). 10.1007/s00439-020-02199-3

30 Solanki, A. K. et al. Mutations in KIRREL1, a slit diaphragm component, cause steroid-resistant nephrotic syndrome. Kidney Int 96, 883–889 (2019). 10.1016/j.kint.2019.06.016

31 Liu, L. et al. Defective nephrin trafficking caused by missense mutations in the NPHS1 gene: insight into the mechanisms of congenital nephrotic syndrome. Hum Mol Genet 10, 2637–2644 (2001).

32 Kitamura, A. et al. A familial childhood-onset relapsing nephrotic syndrome. Kidney Int 71, 946–951 (2007).

33 Shikata, K., Makino, H. & Ota, Z. High resolution three-dimensional meshwork structure of the glomerular basement membrane.Nihon Jinzo Gakkai Shi 34, 379–385 (1992).

34 Khoshnoodi, J., Pedchenko, V. & Hudson, B. G. Mammalian collagen IV. Microsc Res Tech 71, 357–370 (2008). 10.1002/jemt.20564

35 Deen, W. M., Bohrer, M. P. & Brenner, B. M. Macromolecule transport across glomerular capillaries: application of pore theory. Kidney Int 16, 353–365 (1979).

36 Chang, R. S., Robertson, C. R., Deen, W. M. & Brenner, B. M. Permselectivity of the glomerular capillary wall to macromolecules. I. Theoretical considerations. Biophysical Journal 15, 861–886 (1975).

37 Curry, F. E. & Michel, C. C. A fiber matrix model of capillary permeability. Microvasc Res 20, 96–99 (1980).

38 Ogston, A. G. The spaces in a uniform random suspension of fibres. Transactions of the Faraday Society 54, 1754–1757 (1958). 10.1039/TF9585401754

39 Evenson, M. A. & Deutsch, H. F. Influence of fatty acids on the isoelectric point properties of human serum albumin. Clin Chim Acta 89, 341–354 (1978).

40 Kanwar, Y. S. & Farquhar, M. G. Anionic sites in the glomerular basement membrane. In vivo and in vitro localization to the laminae rarae by cationic probes. The Journal of Cell Biology 81, 137–153 (1979).

41 Israelachvili, J. N. Intermolecular and Surface Forces. 3rd Edition edn, Vol. 3rd Edition (Academic Press, 2011).

42 Jeansson, M. & Haraldsson, B. Glomerular size and charge selectivity in the mouse after exposure to glucosaminoglycan-degrading enzymes. J Am Soc Nephrol 14, 1756–1765 (2003).

43 Miner, J. H. & Sanes, J. R. Collagen IV alpha 3, alpha 4, and alpha 5 chains in rodent basal laminae: sequence, distribution, association with laminins, and developmental switches. The Journal of Cell Biology 127, 879–891 (1994).

44 Massoudi, D., Miner, J. H. & Gould, D. B. Collagen IV in Gould syndrome and Alport syndrome. Nat Rev Nephrol 21, 778–793 (2025). 10.1038/s41581-025-00982-x

45 Gunwar, S. et al. Glomerular basement membrane. Identification of a novel disulfide-cross-linked network of alpha3, alpha4, and alpha5 chains of type IV collagen and its implications for the pathogenesis of Alport syndrome. The Journal of Biological Chemistry 273, 8767–8775 (1998).

46 Meehan, D. T. et al. Biomechanical strain causes maladaptive gene regulation, contributing to Alport glomerular disease. Kidney Int 76, 968–976 (2009). 10.1038/ki.2009.324

47 Cosgrove, D. et al. Collagen COL4A3 knockout: a mouse model for autosomal Alport syndrome. Genes Dev 10, 2981–2992 (1996).

48 Case, L. B., Zhang, X., Ditlev, J. A. & Rosen, M. K. Stoichiometry controls activity of phase-separated clusters of actin signaling proteins. Science (New York, N.Y.) 363, 1093–1097 (2019). 10.1126/science.aau6313

49 Huber, T. B. & Benzing, T. The slit diaphragm: a signaling platform to regulate podocyte function. Curr Opin Nephrol Hypertens 14, 211–216 (2005).

50 Hagen, W. J. H., Wan, W. & Briggs, J. A. G. Implementation of a cryo-electron tomography tilt-scheme optimized for high resolution subtomogram averaging. J Struct Biol 197, 191–198 (2017). 10.1016/j.jsb.2016.06.007

51 Zheng, S. Q. et al. MotionCor2: anisotropic correction of beam-induced motion for improved cryo-electron microscopy. Nature Methods 14, 331–332 (2017). 10.1038/nmeth.4193

52 Kremer, J. R., Mastronarde, D. N. & McIntosh, J. R. Computer visualization of three-dimensional image data using IMOD. Journal of Structural Biology 116, 71–76 (1996). DOI 10.1006/jsbi.1996.0013

53 Zheng, S. et al. AreTomo: An integrated software package for automated marker-free, motion-corrected cryo-electron tomographic alignment and reconstruction. J Struct Biol X 6, 100068 (2022). 10.1016/j.yjsbx.2022.100068

54 Tegunov, D. & Cramer, P. Real-time cryo-electron microscopy data preprocessing with Warp. Nature Methods 16, 1146–1152 (2019). 10.1038/s41592-019-0580-y

55 Liu, Y.-T. et al. Isotropic reconstruction for electron tomography with deep learning. Nature Communications 13, 6482 (2022). 10.1038/s41467-022-33957-8

56 Lamm, L. et al. MemBrain v2: an end-to-end tool for the analysis of membranes in cryo-electron tomography. bioRxiv (2024). 10.1101/2024.01.05.574336

57 Pettersen, E. F. et al. UCSF ChimeraX: Structure visualization for researchers, educators, and developers. Protein Sci 30, 70–82 (2021). 10.1002/pro.3943

58 Ermel, U. H., Arghittu, S. M. & Frangakis, A. S. ArtiaX: An electron tomography toolbox for the interactive handling of sub-tomograms in UCSF ChimeraX. Protein Sci 31, e4472 (2022). 10.1002/pro.4472

59 Scheres, S. H. W. RELION: implementation of a Bayesian approach to cryo-EM structure determination. J Struct Biol 180, 519–530 (2012). 10.1016/j.jsb.2012.09.006

60 Nickell, S. et al. TOM software toolbox: acquisition and analysis for electron tomography. J Struct Biol 149, 227–234 (2005).

61 Edelsbrunner, H. & Mücke, E. P. Three-dimensional alpha shapes. ACM Trans. Graph. 13, 43–72 (1994). 10.1145/174462.156635

62 Silverman, B. W. Density Estimation for Statistics and Data Analysis<paper.pdf>. 1st Edition edn, Vol. 1st Edition (Routledge, 1998).

63 McLachlan, G. & Peel, D. Finite Mixture Models. (John Wiley & Sons, Inc., 2000).

64 Schwarz, G. Estimating the Dimension of a Model. The Annals of Statistics 6, 461–464 (1978).

65 Massey Jr, F. J. The Kolmogorov-Smirnov Test for Goodness of Fit. Journal of the American Statistical Association 46, 68–78 (1951). 10.1080/01621459.1951.10500769

66 Bentley, J. L. Multidimensional binary search trees used for associative searching. Commun. ACM 18, 509–517 (1975). 10.1145/361002.361007

67 Schnitzer, J. E. Analysis of steric partition behavior of molecules in membranes using statistical physics. Application to gel chromatography and electrophoresis. Biophysical Journal 54, 1065–1076 (1988).

68 Abdel-Mageed, S. M. & Mohamed, E. I. Comparative modeling of combined transport of water and graded-size molecules across the glomerular capillary wall. J Theor Biol 394, 109–116 (2016). 10.1016/j.jtbi.2016.01.019

