## Supplementary file for "Molecular anatomy of the glomerular filtration barrier reveals topology-encoded albumin sieving"

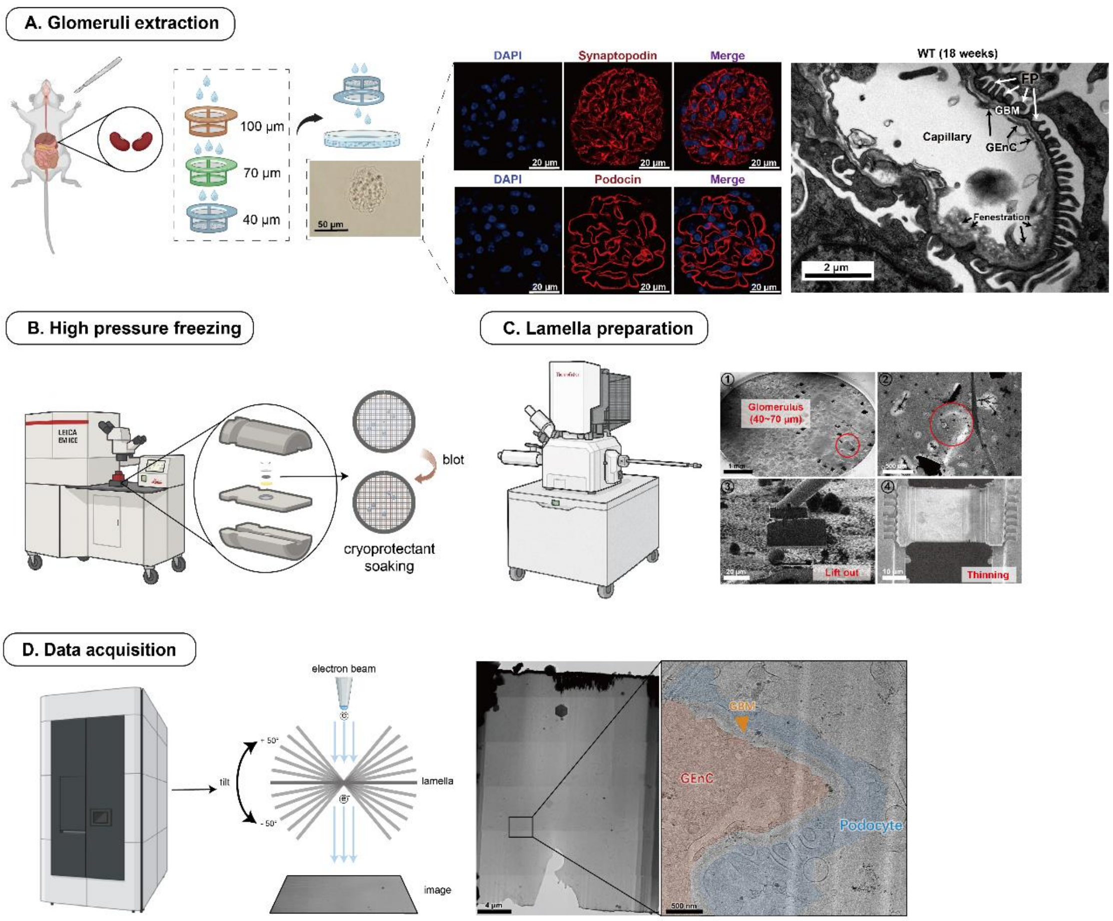


**Extended Data Fig. 1 | Workflow for glomerular sample preparation and cryo-ET imaging**

(A) Isolation and validation of intact glomeruli. Glomeruli were size-selected and confirmed to preserve structural integrity by immunofluorescence and transmission electron microscopy.

(B) High-pressure freezing for cryo-preservation. Isolated glomeruli were rapidly frozen under high pressure to preserve native ultrastructure.

(C) Serial lift-out sectioning. Sequential focused ion beam milling was applied to prepare glomerular lamellae under 200 nm in thickness for cryo-ET imaging.

(D) Cryo-ET data acquisition. Tilt-series images were collected at 300 kV, clearly resolving the three-layered glomerular filtration barrier.

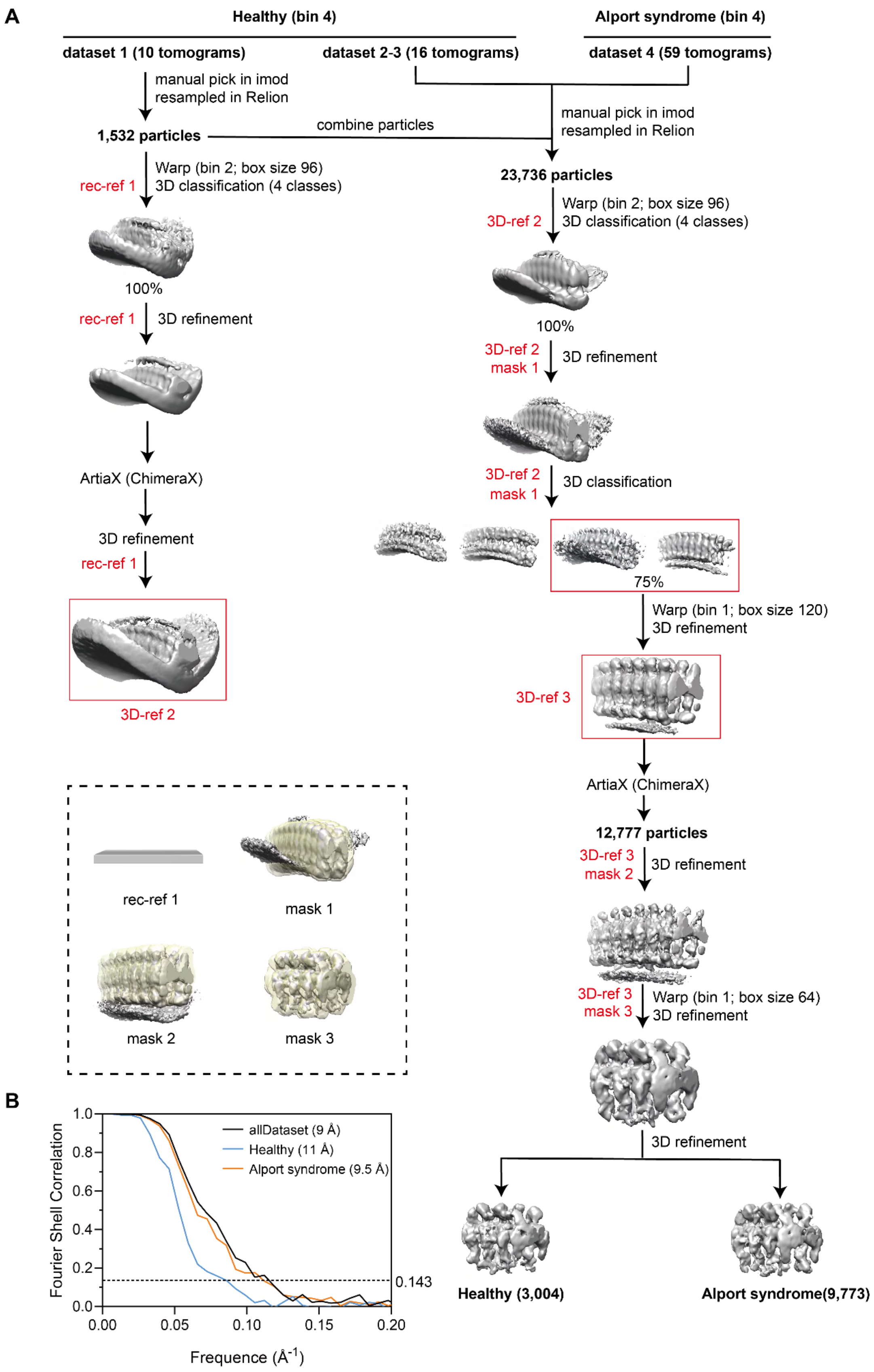


**Extended Data Fig. 2 | Subtomogram averaging of the fenestral rim complex.**

(A) Subtomogram averaging workflow for the fenestral rim complex. Manually picked particles were oversampled at 38 Å based on periodic features. An initial template matching a membrane-like slab (simulating the thickness of plasma membrane) was used for reference. Following 3D classification, particle orientations were manually adjusted in ArtiaX to optimize angular assignments prior to final refinement. The resulting datasets were segregated into two distinct structural maps: Healthy and Alport syndrome.

(B) Resolution validation of the fenestral rim complex. Gold-standard Fourier shell correlation (FSC) curves demonstrate the resolution of the final maps (dashed line: FSC = 0.143 cutoff).

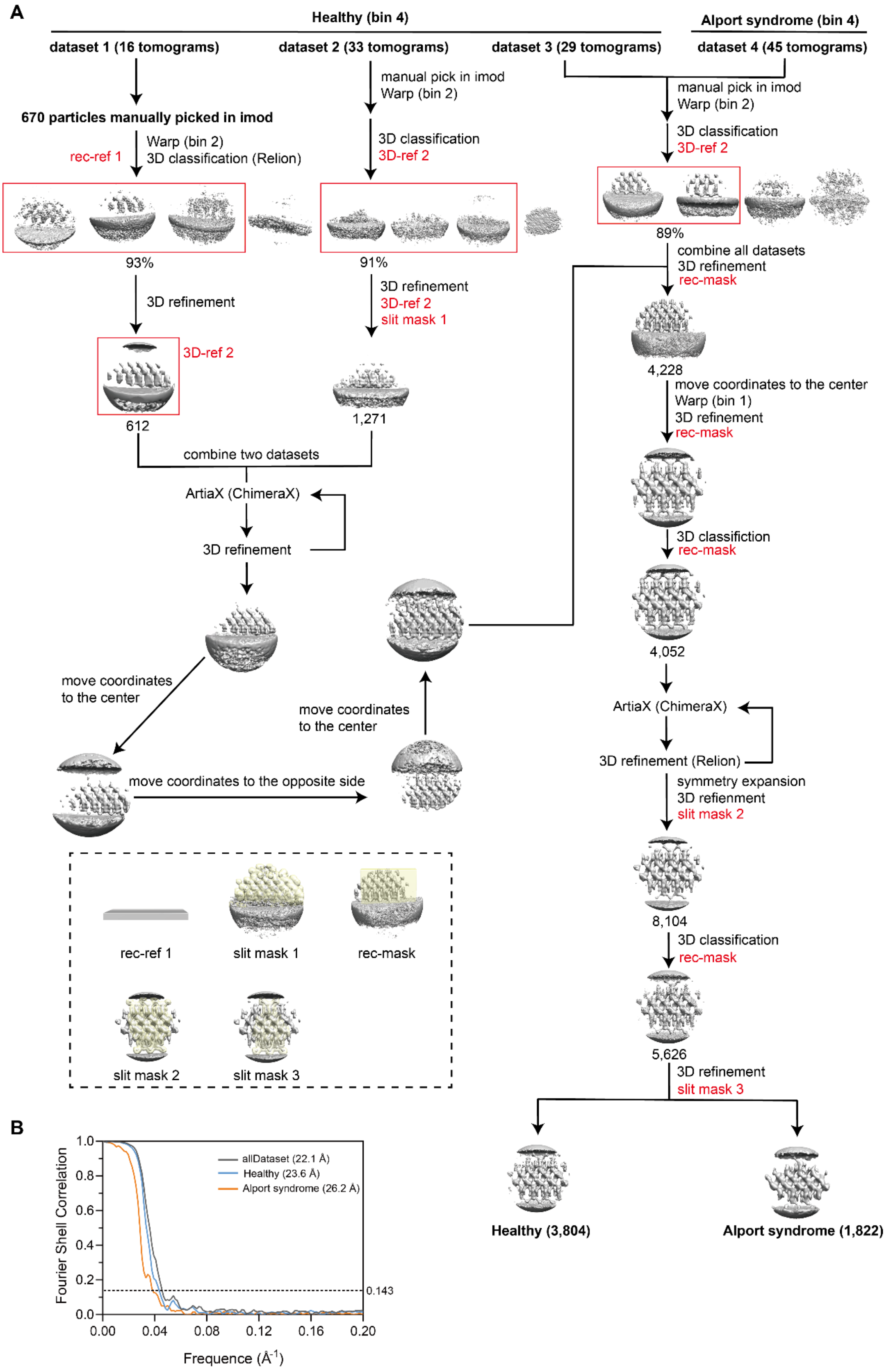


**Extended Data Fig. 3 | Subtomogram averaging of the slit diaphragm.**

(A) Processing pipeline for slit diaphragm subtomogram averaging. Initial particle picking was performed manually, followed by oversampling at 120 Å intervals to account for periodic structural features. A membrane-like slab template, approximating the thickness of plasma membrane, served as the initial reference. After 3D classification, particle orientations were manually adjusted in ArtiaX to improve angular accuracy before final alignment and reconstruction. The processed datasets yielded two distinct structural maps corresponding to Healthy and Alport syndrome conditions.

(B) Global resolution estimates were determined using gold-standard FSC analysis. The final reconstructions reached resolutions where the FSC curves crossed the conventional 0.143 cutoff (dashed line).

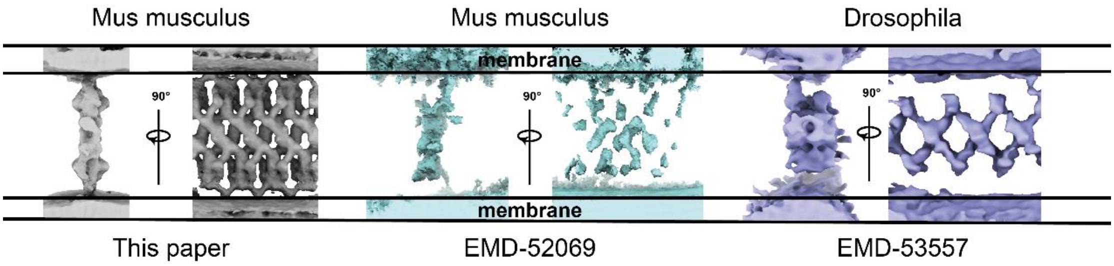


**Extended Data Fig. 4 | Comparison of resolved slit diaphragm subtomogram average maps.**

All slit diaphragm subtomogram average maps are aligned with relative to the plasma membrane. Structures are displayed at the contour levels as suggested in the EMDB: EMD-52069 at 0.00645 and EMD-53557 at 0.32.

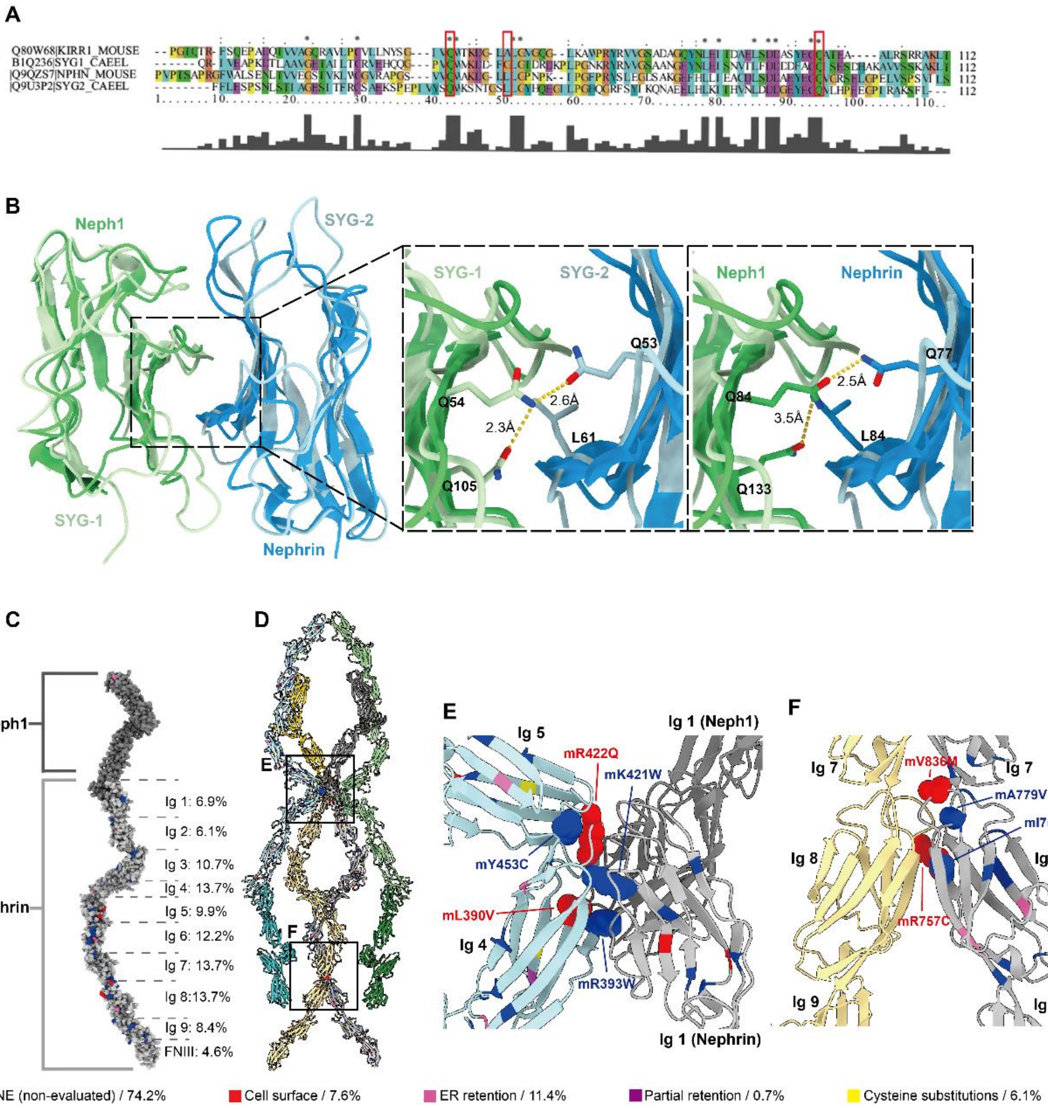


**Extended Data Fig. 5 | Evolutionary conservation, structural homology, and disease-associated mutations in Neph1-Nephrin interaction networks.**

(A) Evolutionary conservation of Neph1 and Nephrin Ig1 domains. Sequence alignment highlights conserved residues between mouse and *C.elegans* (colored by physicochemical properties). Red boxes indicate critical invariant sites.

(B) Structural homology modeling of the Neph1-Nephrin Ig1 interface. Predicted Ig1 domains of Nephrin (AF-Q9QZS7-F1) and Neph1 (AF-Q80W68-F1) were aligned to the SYG-2/SYG-1 heterodimer (PDB: 4OFY). Insets show corresponding atomic interactions between SYG-1/Neph1 (light green/green) and SYG-2/Nephrin (light blue/blue).

(C) Spatial mapping of disease-associated mutations. Clinically identified mutations (n = 131) were mapped onto the heterodimer model and color-coded by functional domains. Numeric labels indicate the percentage of mutations localized to each domain of Nephrin.

(D) Mutational hotspots within the modeled complex. Disease-associated mutations are distributed across oligomeric interfaces, with black boxes highlighting residues predicted to destabilize the complex.

(E) Close-up view of the Neph1(Ig1)-Nephrin (Ig1)-Nephrin (Ig4/5) interaction interfaces. Disease-associated mutation sites in spatial proximity are represented as spheres. Variants reported to retain plasma membrane localization are colored red, suggesting that their pathogenicity arises from impaired complex assembly despite correct trafficking.

(F) Nephrin (Ig8) homodimer interface. Sphere labels indicate mutation sites likely perturbing Ig8-Ig8 interactions.

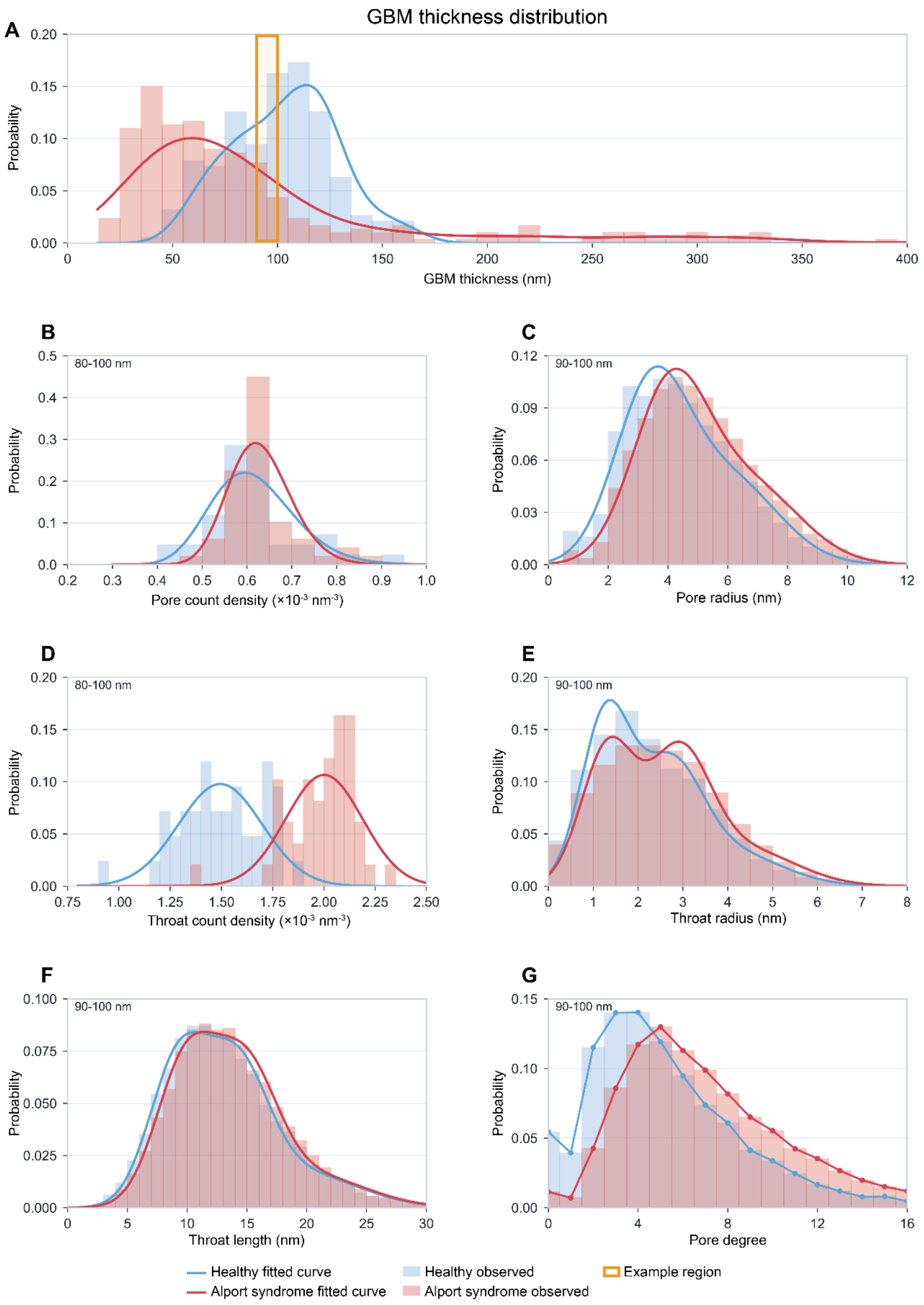


**Extended Data Fig. 6 | Thickness distribution and structural descriptor distributions example from real GBM subsample.**

(A) Thickness distributions of Healthy and Alport syndrome GBM subsamples, with the representative 90–100 nm thickness interval selected as a representative interval.

(B–G) Observed real-sample distributions and corresponding fitted distributions for pore count density, pore radius, throat count density, throat radius, throat length, and pore degree. For density descriptors, the corresponding 80–100 nm interval is shown; other descriptors are shown for the 90–100 nm interval. Pore degree was represented by an empirical probability mass function. Blue indicates Healthy and red indicates Alport syndrome. The full statistical fitting and parameterization workflow is described in Methods. Full fitting results and source data are provided in the GitHub repository.

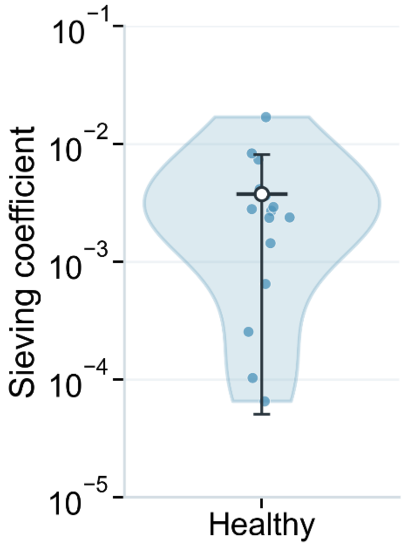


**Extended Data Fig. 7 | Distribution of albumin sieving coefficients in Healthy GBM samples.**

Violin plot shows predicted positive albumin sieving coefficients calculated from the pore-throat transport model for real Healthy large-samples. Each blue dot represents one Healthy sample with a connected albumin-accessible pathway (n = 15). The white point and error bar indicate the arithmetic mean ± standard deviation of the positive values (0.004 ± 0.004), with values ranging from 0.000065 to 0.0170. The remaining 5 Healthy samples have no connected albumin-accessible pathway and are not displayed on the log-scale violin plot; when included as zero sieving coefficients, the Healthy arithmetic mean is 0.003.

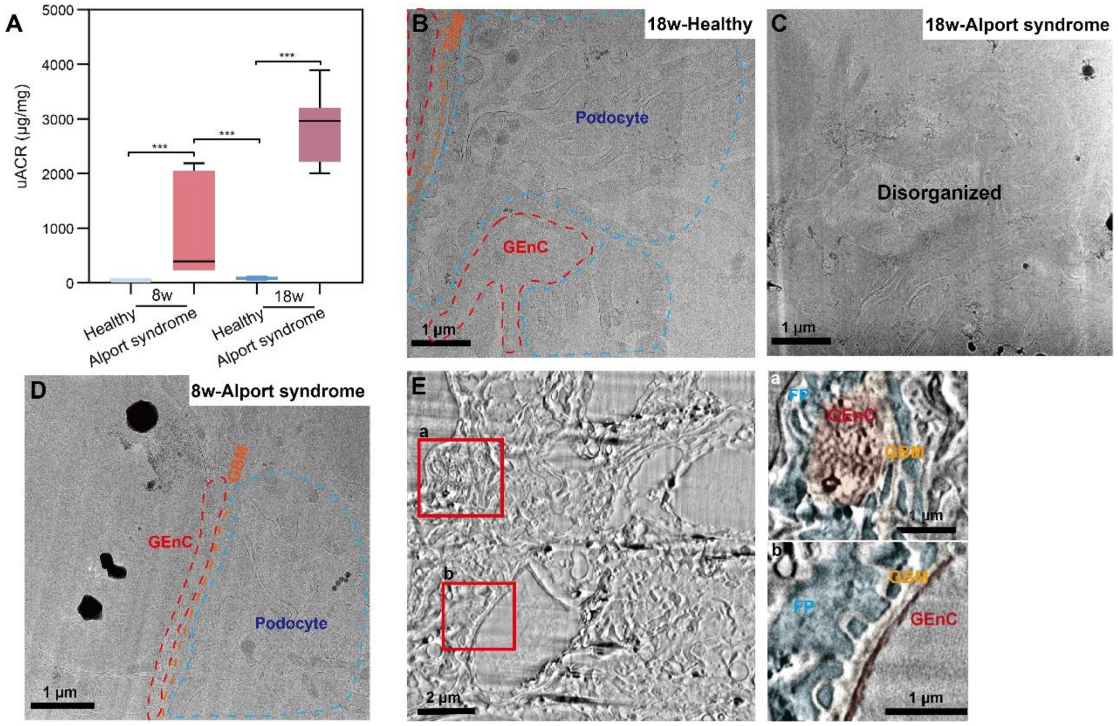


**Extended Data Fig. 8 | Establishment of an early-stage Alport syndrome model with preserved glomerular filtration barrier ultrastructure.**

(A) Urinary albumin-to-creatinine ratio (uACR, μg/mg) in Healthy and Alport syndrome mice at 8 and 18 weeks of age. Mean values: 8w-Healthy, 43.6 μg/mg (n=7); 8w-Alport syndrome, 949.7 μg/mg (n=7); 18w-Healthy, 85.5 μg/mg (n=7); 18w-Alport syndrome, 2893.7 μg/mg (n=7). Statistical comparisons were performed using Mann-Whitney U test for 8w-Healthy versus 8w-Alport syndrome and 8w-Alport syndrome versus 18w Healthy; unpaired student's t-test for 18w Healthy vs. 18w Alport syndrome. P-values were adjusted using the Bonferroni correction for multiple comparisons. Significance: *$p < 0.05$, **$p < 0.01$, ***$p < 0.001$; ns = not significant.

(B) A cryo-TEM image of a glomerulus from an 18-week-old Healthy mouse. The GEnC (red dashed line), GBM (orange dashed line) and podocyte foot processes (blue dashed line) are demarcated to illustrate the tripartite glomerular filtration barrier (GFB).

(C) A cryo-TEM image of a glomerulus from an 18-week-old Alport syndrome mouse. At this stage, the structural integrity of the GFB is severely compromised, rendering the distinct layers of the GEnC, GBM and podocytes indistinguishable.

(D) A cryo-TEM image of a glomerulus from an 8-week-old Alport syndrome mouse. The GEnC,

GBM and podocyte foot processes (blue dashed line) remain morphologically discernible, indicating early-stage pathology.

(E) Cryo-PFIB/SEM volume image of the GFB in an 8-week-old Alport syndrome mouse, showing preserved native ultrastructure. Red boxes highlight two GFB regions (a and b) shown in different views. Color-coded structures: Podocyte foot processes (blue); GBM (orange); GEnC (red).

Scale bars: 1 μm in B–D and insets a, b of E; 2 μm in E.

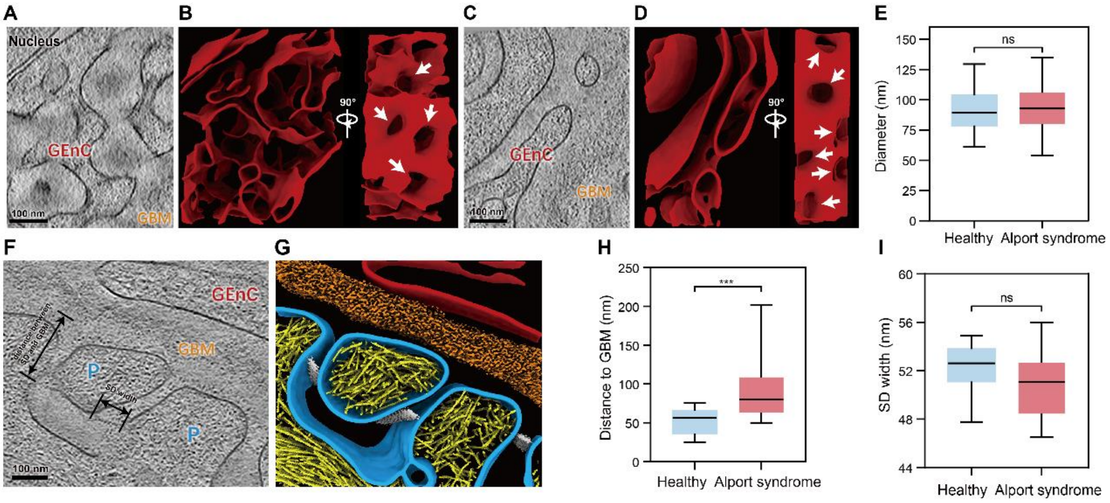


**Extended Data Fig. 9 | Comparative analysis of endothelial fenestrations and slit diaphragm architecture in Healthy and Alport syndrome mice.**

(A–B) Representative tomographic cross-section of a GEnC from a Healthy mouse (A) and the corresponding 3D surface renderings in cross-sectional (left) and membrane (right) views (B). White arrows indicate fenestrations.

(C–D) Representative tomographic cross-section of a GEnC from an Alport syndrome mouse (C) and the corresponding 3D surface renderings (D). White arrows denote fenestrations.

(E) Comparison of fenestral diameters between Healthy and Alport syndrome mice. The mean diameters are 91.3 nm (n = 100) and 93.6 nm (n = 100) for Healthy and Alport syndrome, respectively. Statistical significance was determined using a Mann-Whitney U test (ns: not significant; $*p < 0.05$, $**p < 0.01$, $***p < 0.001$).

(F–G) Ultrastructural organization of the glomerular filtration barrier in Alport syndrome mice, shown as a representative tomographic slice (F) and the corresponding 3D segmentation (G). Black lines indicate measured parameters in (F).

(H) Quantification of the distance between the slit diaphragm (SD) and the GBM. Mean distances are 53.2 nm (Healthy; n = 30) and 90.8 nm (Alport syndrome; n = 30). Statistical significance was determined using a Mann-Whitney U test.

(I) Quantification of slit diaphragm width. Mean widths are 52.0 nm (Healthy; n = 25) and 50.9 nm (Alport syndrome; n = 25). Statistical significance was determined using unpaired student's t-test.

Scale bars: 100 nm in A, C and F.

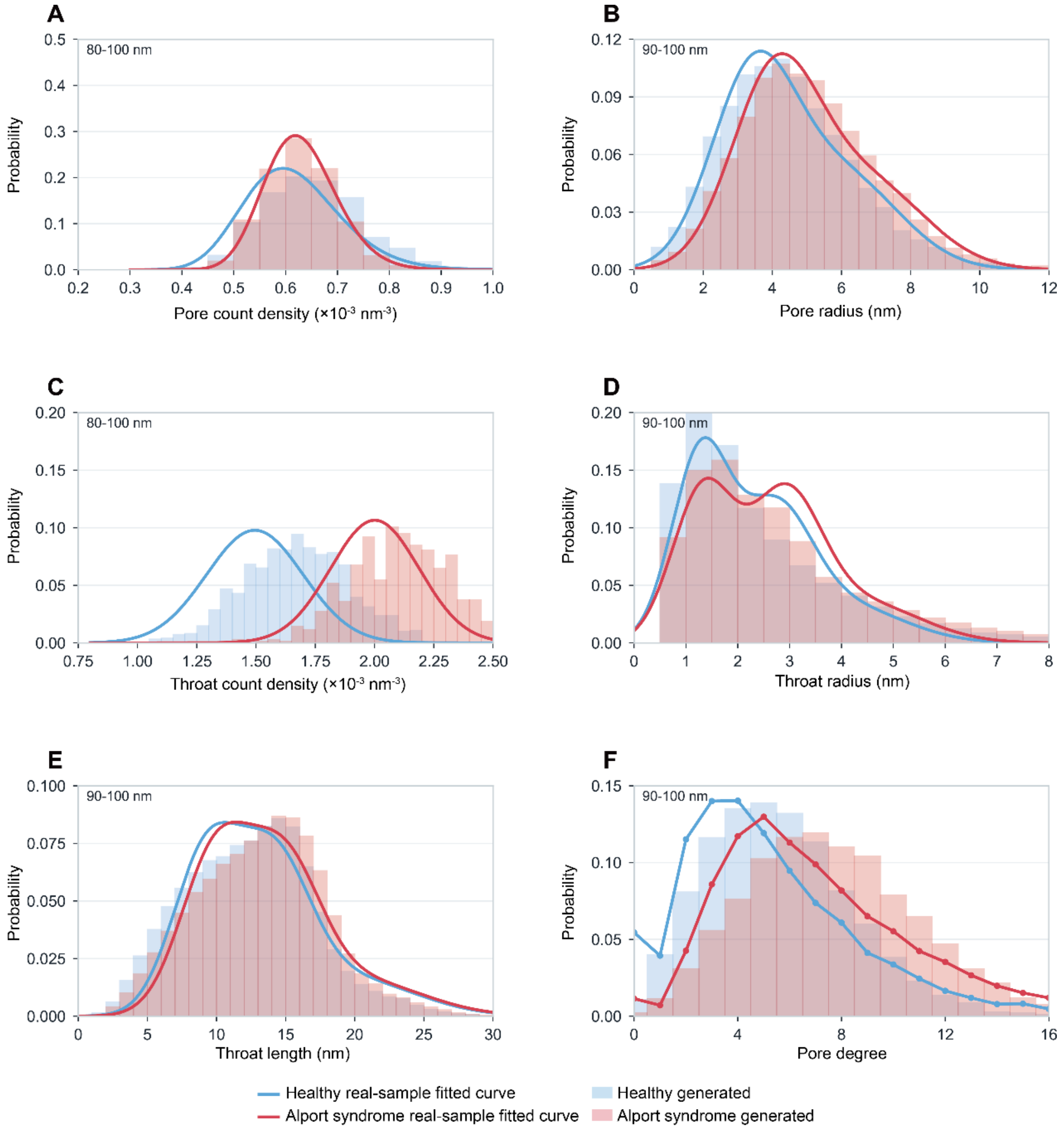


**Extended Data Fig. 10 | Generated-network descriptor distribution examples compared with real-sample fitted distributions.**

(A–F) Distributions measured from generated GBM pore-throat networks for pore count density, pore radius, throat count density, throat radius, throat length, and pore degree in the same representative thickness interval of Extended Data Fig.6. Bars show generated-network statistics, whereas solid curves show the real-sample fitted target distributions or empirical probability mass functions used for network generation. Density descriptors are shown for the 80–100 nm bin, and object-level descriptors are shown for the 90–100 nm bin. Blue indicates Healthy and red indicates Alport syndrome. Full fitting results and source data are provided in the GitHub repository.

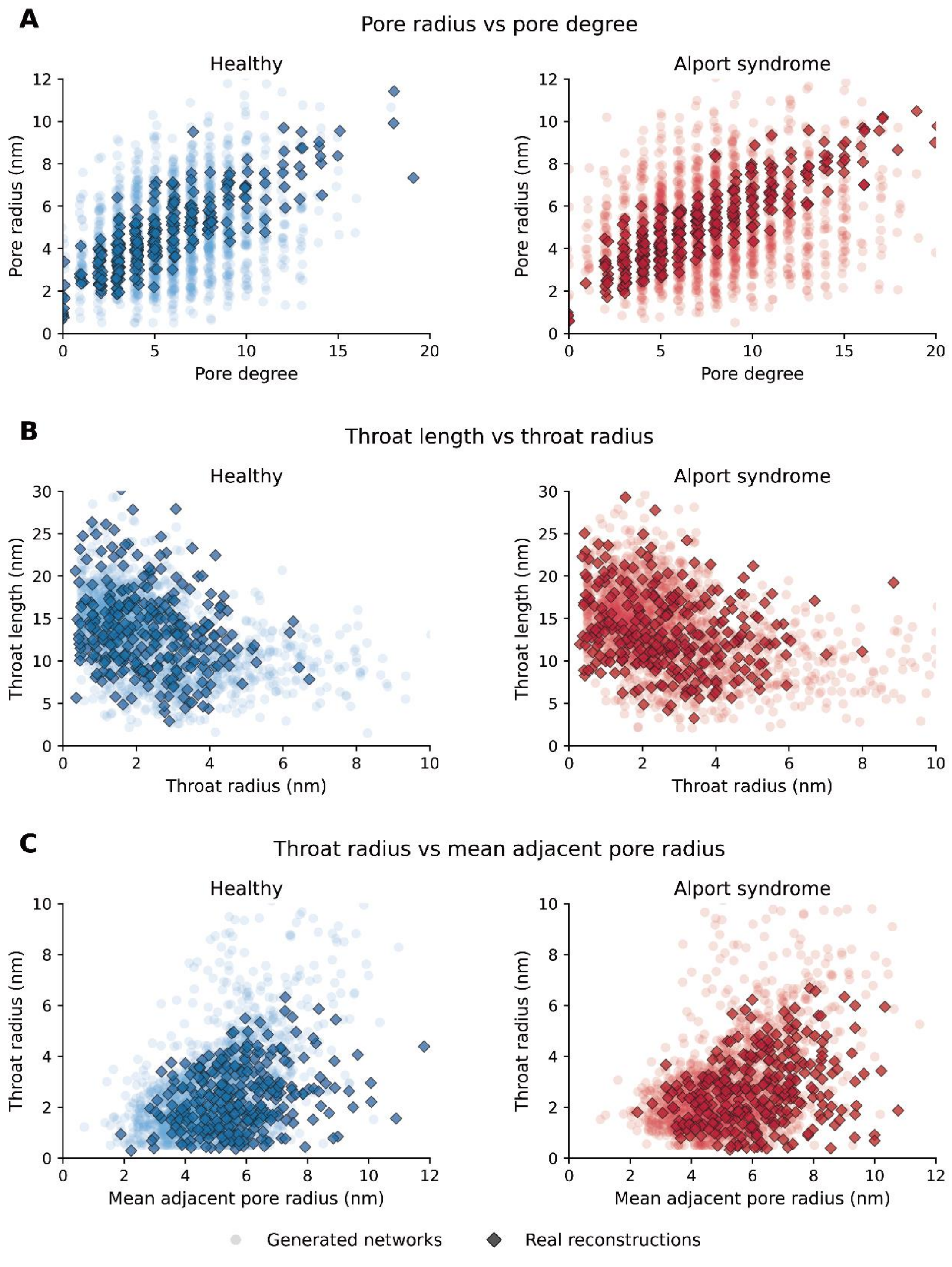


**Extended Data Fig. 11 | Structural-descriptor correlations in GBM networks.**

Each panel compares randomly sampled measurements from generated GBM networks with measurements from real reconstructed GBM subsamples. For visualization clarity, 1,500 generated measurements and 300 real reconstructed measurements were randomly sampled for each group in each descriptor pair. Light blue and light red circles indicate generated Healthy and Alport syndrome networks, respectively, and dark diamond markers indicate real reconstructed measurements from the corresponding group.

(A) Pore radius versus pore degree.

(B) Throat length versus throat radius.

(C) Throat radius versus mean adjacent pore radius.

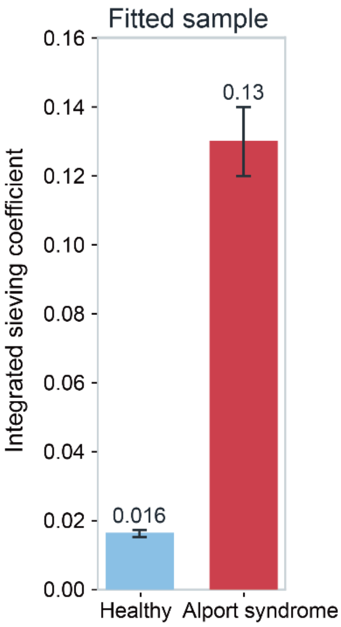


**Extended Data Fig. 12 | Integrated fitted GBM sieving coefficient comparison.**

Integrated GBM-level albumin sieving coefficients calculated from generated networks. Values are integrated coefficient ± bootstrap standard deviation: 0.016 ± 0.001 for Healthy and 0.13 ± 0.01 for Alport syndrome. Standard deviations were calculated from 200 thickness-stratified bootstrap ensembles of 1,500 networks per group.

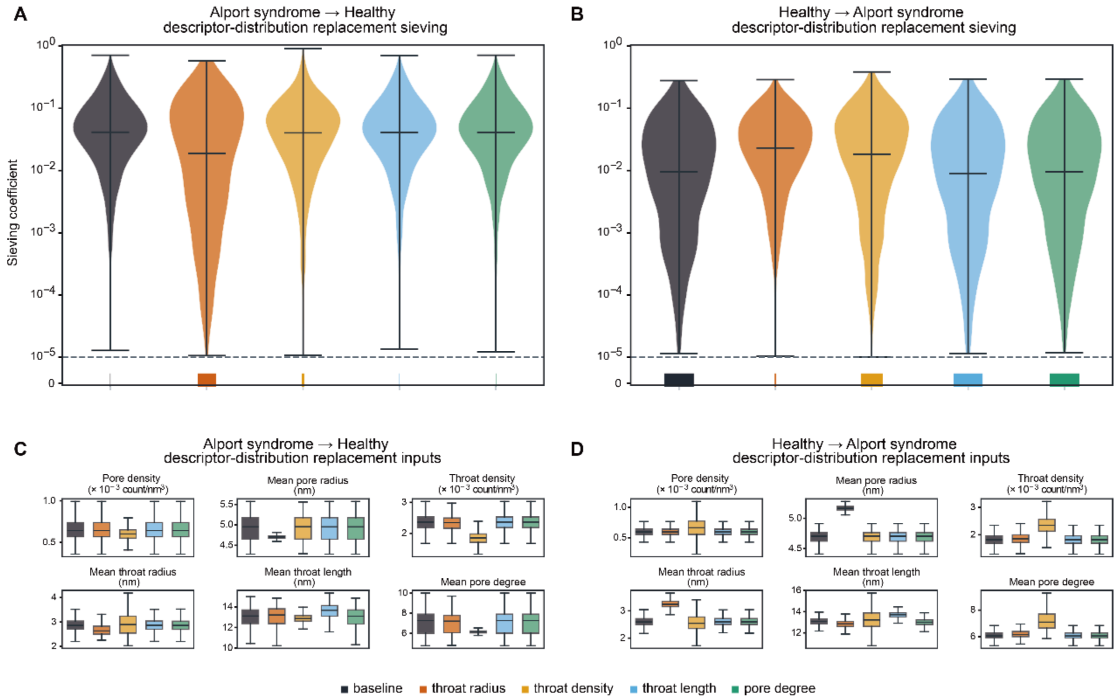


**Extended Data Fig. 13 | Descriptor-distribution replacement analysis of GBM sieving.**

(A–B) Albumin sieving coefficients after single descriptor-distribution replacement. In (A), Alport syndrome GBM networks were generated with Alport descriptor distributions replaced by the corresponding Healthy distributions. In (B), Healthy GBM networks were generated with Healthy descriptor distributions replaced by the corresponding Alport syndrome distributions. Baseline indicates networks generated using the original condition-specific descriptor distributions without replacement. Colors denote baseline, throat-radius replacement, throat-density replacement, throat-length replacement, and pore-degree replacement. Violin plots show sieving coefficients at or above $10^{-5}$ on a logarithmic scale; the dashed horizontal line marks the plotting threshold at $10^{-5}$, and median lines are shown within each violin. Colored bars at zero indicate generated networks with zero sieving coefficients or coefficients below $10^{-5}$. Mean per-network sieving coefficients and zero-row fractions are reported in Extended Data Table 2A.

(C–D) Generated structural-descriptor distributions under the same baseline and replacement conditions. Boxplots summarize pore density, mean pore radius, throat density, mean throat radius, mean throat length, and mean pore degree in the generated networks. Boxes indicate median and interquartile range (IQR), whiskers indicate the 1.5 × IQR, and outliers are not shown. Numerical summaries are reported as median [1.5 × IQR whisker range] in Extended Data Table 2B.

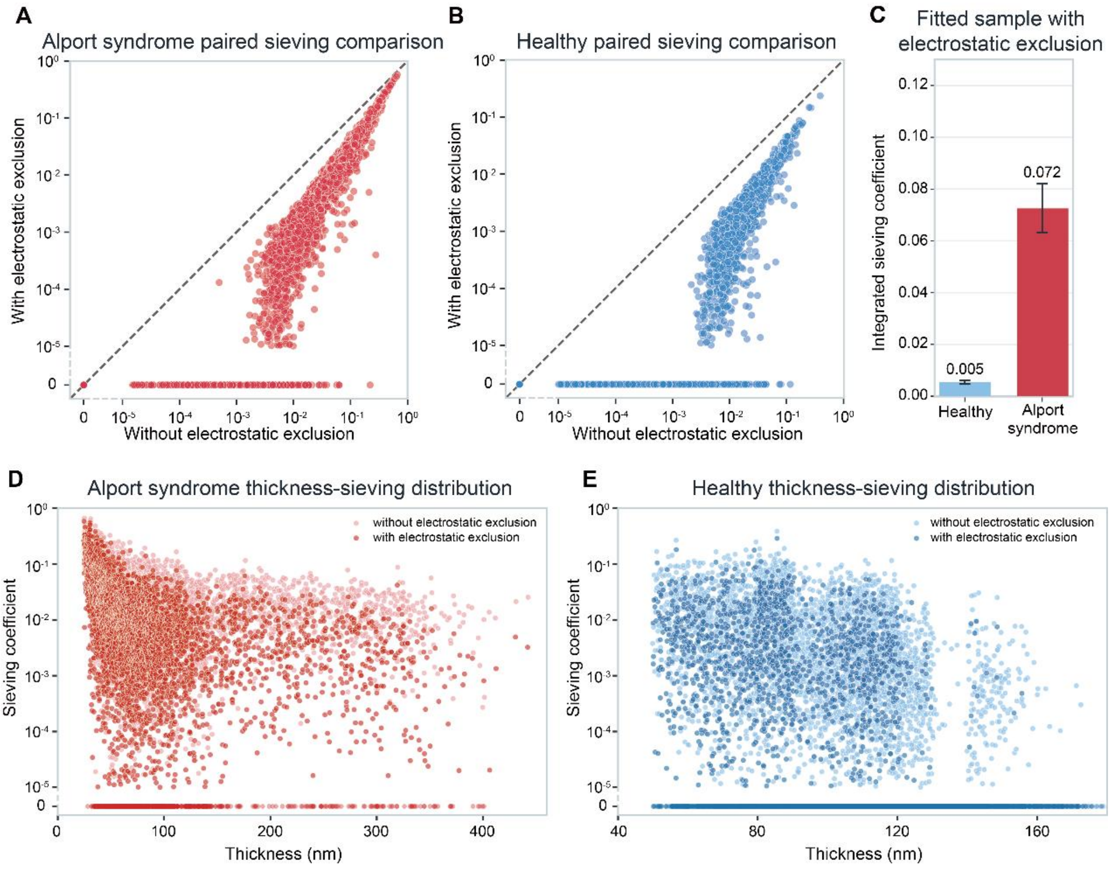


**Extended Data Fig. 14 | Effect of electrostatic exclusion on generated-network sieving.** Albumin sieving was recalculated in the same generated GBM networks after increasing the albumin effective radius from 3.55 nm to 4.25 nm, as an effective-radius approximation for electrostatic exclusion.

(A–B) Paired comparison of albumin sieving coefficients in the same Alport syndrome (A) and Healthy (B) generated networks before and after applying electrostatic exclusion through an increased albumin effective radius. The x-axis shows sieving without electrostatic exclusion, and the y-axis shows sieving with electrostatic exclusion. Points on the zero line indicate networks with sieving coefficients below $10^{-5}$.

(C) Integrated GBM-level albumin sieving coefficients after applying electrostatic exclusion. Values are integrated coefficient ± bootstrap standard deviation: 0.0050 ± 0.0006 for Healthy and 0.072 ± 0.009 for Alport syndrome. Standard deviations were calculated from 200 thickness-stratified bootstrap ensembles of 1,500 networks per group.

(D–E) Thickness-sieving distributions for Alport syndrome (D) and Healthy (E) generated networks at the two effective radii. At both solute radii, all 10,000 generated networks yielded valid calculations, including 5,000 Healthy and 5,000 Alport syndrome networks. For visualization, networks with either zero sieving or sieving coefficients below the plotting threshold of $10^{-5}$ were displayed at the zero row. The corresponding fraction of networks with sieving coefficients above the plotting threshold decreased from 74.6% to 28.9% in Healthy networks and from 99.6% to 85.0% in Alport syndrome networks.

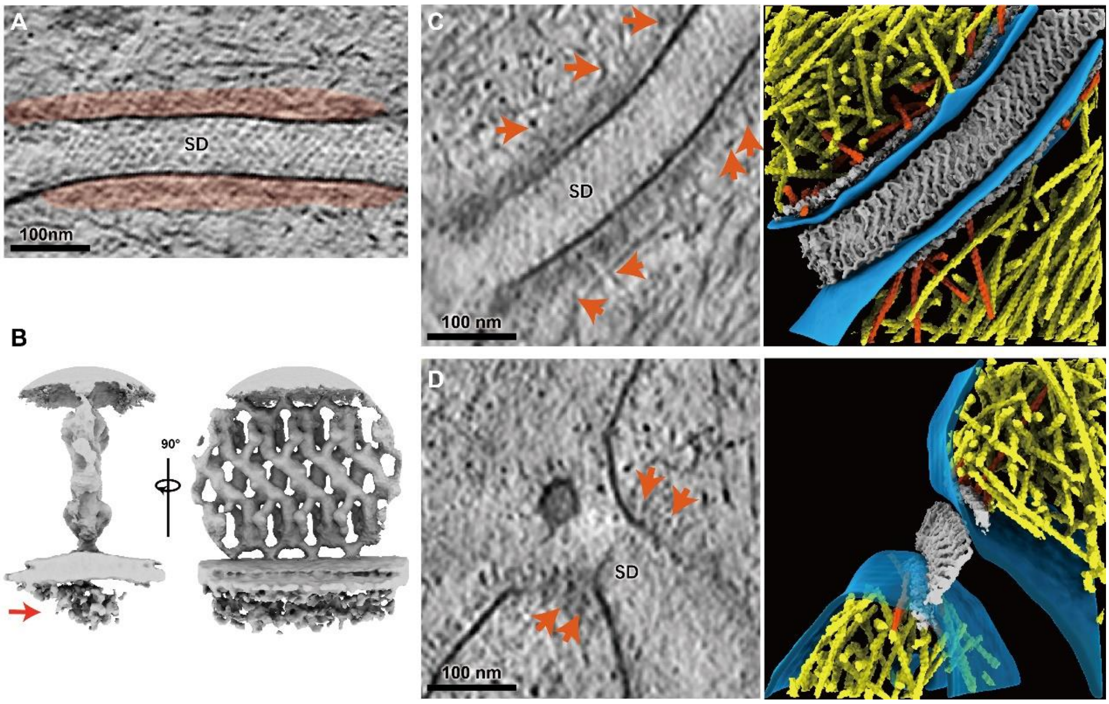


**Extended Data Fig. 15 | Cytosolic density beneath the slit diaphragm and its association with the actin cytoskeleton.**

(A) An unidentified density is observed on the cytoplasmic side of the slit diaphragm (SD). The red translucent overlay highlights this dense region.

(B) Subtomogram averaged map revealing the architecture of the slit diaphragm cytosolic density, indicated by red arrow.

(C–D) Association between actin filaments and the slit diaphragm cytosolic density, presented in two orthogonal views. Left panels: representative tomographic slices; right panels: corresponding 3D renderings. Actin filaments in close proximity to th slit diaphragm cytosolic density are highlighted in orange.

Scale bars: 100 nm in A, C and D.

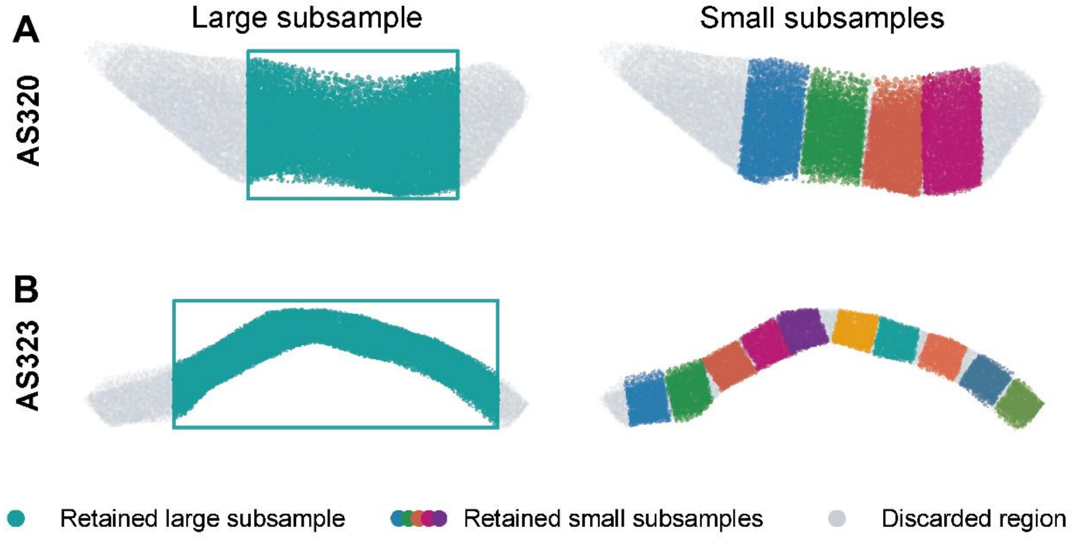


**Extended Data Fig. 16 | Large- and small-subsample selection example.** Representative Alport syndrome samples illustrating retained large subsamples and their corresponding small subsamples. Teal points denote the retained large subsample, multicolored points denote retained small subsamples, and gray points denote discarded regions.

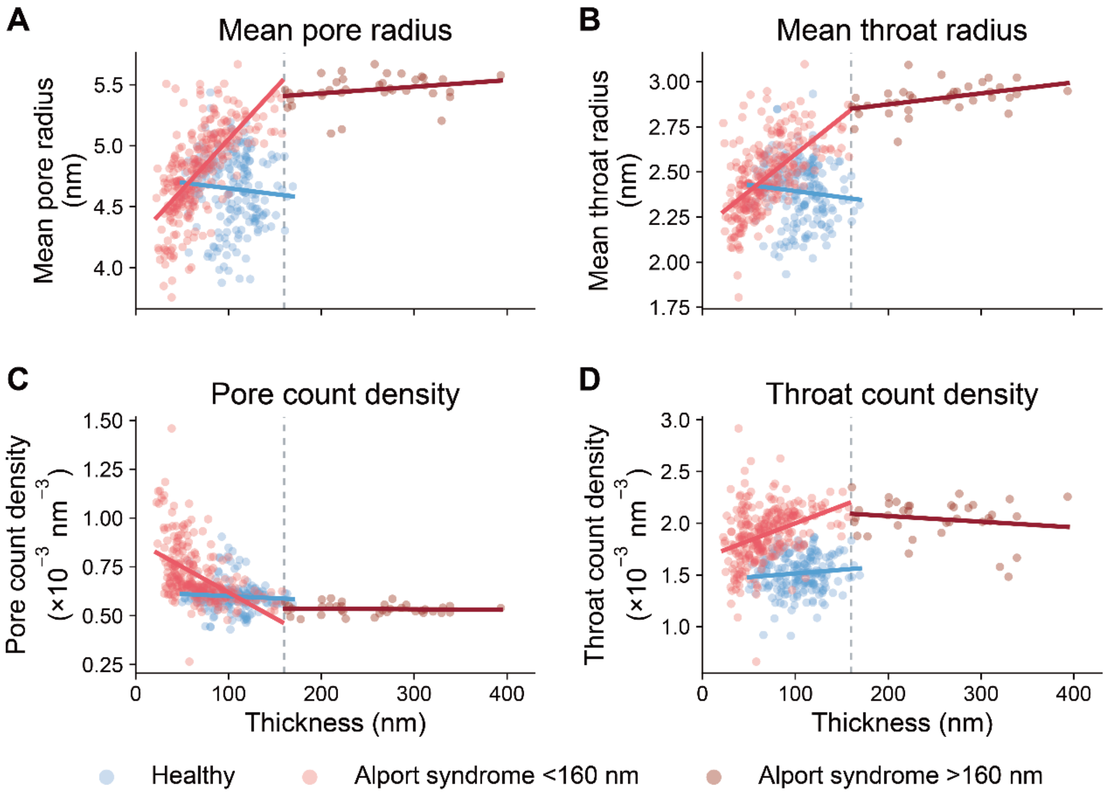


**Extended Data Fig. 17 | Thickness-dependent structural descriptors in Healthy and Alport syndrome subsamples.**

Scatter plots show subsample-level relationships between GBM thickness and mean pore radius, mean throat radius, pore count density, and throat count density. Blue points indicate Healthy subsamples, light red points indicate Alport syndrome subsamples with thickness <160 nm, and dark red points indicate Alport syndrome subsamples with thickness >160 nm. Solid lines show group-wise linear regression fits, and the dashed vertical line marks 160 nm. Regression slopes are reported as changes per 100 nm increase in GBM thickness in the order Healthy, Alport syndrome <160 nm, and Alport syndrome >160 nm: mean pore radius (–0.095, 0.828, 0.054 nm), mean throat radius (–0.068, 0.411, 0.061 nm), pore count density (–0.021, –0.263, –0.003 ×10$^{-3}$ nm$^{-3}$), and throat count density (0.072, 0.339, –0.055 ×10$^{-3}$ nm$^{-3}$).

**Extended Data Table 1. Parameters used in albumin transport model**

| Parameter | Symbol | Value | Unit | Source |
|---|---|---|---|---|
| Dynamic viscosity of water at 37°C | $\mu$ | $6.91 \times 10^{-4}$ | Pa·s | [69] |
| Free diffusion coefficient of albumin at 37°C | $D_{alb}$ | $9.26 \times 10^{-11}$ | m²/s | Calculated from the Stokes-Einstein relation; [70] |
| Baseline albumin hydrodynamic radius | $r_{\mathrm{alb}}$ | 3.55 | nm | [1] |
| Bjerrum length in water | $l_B$ | 0.7 | nm | Calculated from $l_B = e^2/(4\pi\varepsilon_0\varepsilon_r k_B T)$; [41] |
| Capillary hydrostatic pressure | $P_{\mathrm{GC}}$ | 55 | mmHg | [71] |
| Plasma oncotic pressure | $\pi_{\mathrm{GC}}$ | 30 | mmHg | [71] |
| Bowman's space hydrostatic pressure | $P_{\mathrm{BS}}$ | 15 | mmHg | [71] |
| Generated network in-plane size | $L \times L$ | $200 \times 200$ | nm² | Model setting |
| Bulk-layer thickness | $L_{\mathrm{bulk}}$ | 30 | nm | Estimated from structure |
| Bulk-layer cross-sectional area | $A_{\mathrm{bulk}}$ | 16000 | nm² | Model setting |
| Slit diaphragm effective diffusion length | $L_{\mathrm{SD}}$ | 10 | nm | Estimated from structure |
| Slit diaphragm water-accessible area | $A_{\mathrm{SD}}$ | 5400 | nm² | Estimated from structure |
| Fiber radius in the stochastic fiber-network model | $r_f$ | 1.5 | nm | Estimated from structure |
| Fiber volume fraction in the stochastic fiber-network model | $V_f$ | 0.55 | - | Estimated from structure |

**Extended Data Table 2. Numerical summary of descriptor-distribution replacement analysis.**

**A. Per-network sieving summary**

| Direction | Condition | Mean sieving coefficient | Zero-row fraction |
|---|---|---|---|
| Alport syndrome → Healthy | Baseline | 0.0685 | 0.9% |
| Alport syndrome→ Healthy | Throat radius | 0.0456 | 24.5% |
| Alport syndrome → Healthy | Throat density | 0.0634 | 3.8% |
| Alport syndrome → Healthy | Throat length | 0.0677 | 0.8% |
| Alport syndrome → Healthy | Pore degree | 0.0684 | 0.8% |
| Healthy → Alport syndrome | Baseline | 0.0171 | 39.4% |
| Healthy → Alport syndrome | Throat radius | 0.0379 | 1.5% |
| Healthy → Alport syndrome | Throat density | 0.0276 | 29.0% |
| Healthy → Alport syndrome | Throat length | 0.0167 | 38.8% |
| Healthy → Alport syndrome | Pore degree | 0.0172 | 39.0% |

**B. Generated structural descriptors**

Values are median [1.5×IQR whisker range]. Pore and throat densities are shown as $\times 10^{-3}\text{nm}^{-3}$); radii and lengths are in nm.

| Direction | Condition | Pore density | Pore radius | Throat density | Throat radius | Throat length | Pore degree |
|---|---|---|---|---|---|---|---|
| Alport syndrome→ Healthy | Baseline | 0.643 [0.359–0.993] | 4.950 [4.283–5.571] | 2.363 [1.670–3.032] | 2.863 [2.203–3.524] | 13.122 [10.447–14.993] | 7.277 [4.784–10.009] |

| Direction | Condition | Pore density | Pore radius | Throat density | Throat radius | Throat length | Pore degree |
|---|---|---|---|---|---|---|---|
| Alport syndrome → Healthy | Throat radius | 0.643 [0.359–0.993] | 4.702 [4.592–4.815] | 2.341 [1.670–2.976] | 2.636 [2.259–3.318] | 13.219 [10.229–14.849] | 7.220 [4.810–9.712] |
| Alport syndrome → Healthy | Throat density | 0.602 [0.409–0.796] | 4.953 [4.293–5.559] | 1.849 [1.325–2.389] | 2.894 [2.032–4.187] | 12.866 [11.847–13.970] | 6.149 [5.804–6.538] |
| Alport syndrome → Healthy | Throat length | 0.643 [0.359–0.993] | 4.950 [4.283–5.571] | 2.363 [1.670–3.032] | 2.868 [2.208–3.521] | 13.678 [11.570–15.347] | 7.277 [4.784–10.009] |
| Alport syndrome → Healthy | Pore degree | 0.643 [0.359–0.993] | 4.950 [4.283–5.571] | 2.363 [1.670–3.032] | 2.862 [2.203–3.532] | 13.092 [10.322–14.848] | 7.277 [4.784–10.009] |
| Healthy →Alport syndrome | Baseline | 0.596 [0.427–0.765] | 4.706 [4.411–4.912] | 1.825 [1.293–2.352] | 2.594 [2.172–3.046] | 13.076 [12.175–13.962] | 6.061 [5.323–6.837] |
| Healthy →Alport syndrome | Throat radius | 0.596 [0.427–0.765] | 5.166 [5.056–5.278] | 1.858 [1.307–2.413] | 3.244 [2.864–3.657] | 12.850 [11.891–13.784] | 6.170 [5.448–6.959] |
| Healthy →Alport syndrome | Throat density | 0.663 [0.231–1.095] | 4.707 [4.410–4.911] | 2.349 [1.531–3.221] | 2.545 [1.721–3.410] | 13.218 [10.784–15.761] | 7.095 [5.858–9.281] |
| Healthy →Alport syndrome | Throat length | 0.596 [0.427–0.765] | 4.706 [4.411–4.912] | 1.825 [1.293–2.352] | 2.599 [2.194–3.040] | 13.719 [12.919–14.459] | 6.061 [5.323–6.837] |
| Healthy →Alport syndrome | Pore degree | 0.596 [0.427–0.765] | 4.706 [4.411–4.912] | 1.825 [1.293–2.352] | 2.597 [2.179–3.044] | 13.001 [12.107–13.898] | 6.061 [5.323–6.837] |

**Extended Data Table 3. Principal assumptions of albumin-transport model**

| Model component | Assumption |
|---|---|
| General transport | Albumin transport was assumed to be steady and one-dimensional within each transport element. |
| Albumin conservation | Albumin was assumed to be conserved, with no adsorption or degradation within the filtration pathway. |
| SD representation | The slit diaphragm was represented as a single transport layer, with each major lattice opening approximated as a deformable hexagonal pore whose geometry varied with slit diaphragm width. |
| SD albumin geometry | Albumin was approximated as a triaxial ellipsoid. |
| SD mixing | The region downstream of the slit diaphragm was assumed to be a well-mixed reservoir. |
| GBM representation | The albumin-selective region of the GBM was represented by the pore-throat network extracted from the lamina densa. The throats were approximated as cylindrical channels. |
| GBM mixing | Each GBM pore was represented by a single equivalent albumin concentration, and the common downstream reservoir was assumed to be well mixed. |
| GBM albumin geometry | Albumin was approximated as a sphere. |
| Electrostatic exclusion | Electrostatic repulsion was approximated as an additional geometric exclusion distance, while the hydrodynamic transport properties of hydrated albumin were assumed to remain unchanged. |
| Bulk-fluid regions | The bulk-fluid regions were assumed to impose no geometric sieving and were represented as one-dimensional advection-diffusion layers. |
